# Lower Airway Dysbiosis in NTM+ Bronchiectasis is Associated with NET-Predominant Severe Phenotypes

**DOI:** 10.1101/2025.09.18.677189

**Authors:** Shivani Singh, Fares Darawshy, Kirby Erlandson, Jayanth Kumar Narayana, Qingsheng Li, Yonghua Li, Isabella Atandi, Kelsey Krolikowski, Shrey Patel, Destiny Collazo, Micheál Mac Aogáin, Amy Gilmour, Merete Long, Miao Chang, Afshana Hoque, Rosemary Schluger, Sanjan Kumar, Cecilia J. Chung, Kendrew Wong, Gabriella Porter, Yicheng Feng, Anna Czachor, Colin McCormick, Emily Clementi, Yaa Kyeremateng, Alena Lukovnikova, Danielle Harris, Sebastian Gomez, Taylor Kain, Ibrahim Kocak, Rajbir Singh, Claudia Rodriguez, Benjamin Kwok, Clea Barnett, Matthias Kugler, Michael D. Weiden, Nathaniel Nelson, Jake G. Natalini, David Luglio, Ludovic Desvignes, Samir Gautam, Erin McGuire, Terry Gordon, Imran Sulaiman, Jun-Chieh J. Tsay, Ashwin Basavaraj, Benjamin G. Wu, David Kamelhar, Doreen Addrizzo-Harris, James D. Chalmers, Sanjay H. Chotirmall, Leopoldo N. Segal

## Abstract

**Rationale:** The discoveries of neutrophilic inflammation and *Pseudomonas*-dominant pulmonary dysbiosis have helped pave the way for host-directed therapy in bronchiectasis. Substantial knowledge gaps remain about the interplay between neutrophilic signatures and microbes in non-tuberculous mycobacterial lung disease (NTM-LD), a phenotypically diverse lung infection that is increasingly prevalent in the United States and other parts of the world.

**Objectives:** Evaluate the lower airway microbiota and neutrophilic traits in NTM- and NTM+ bronchiectasis.

**Methods:** 16S rRNA gene sequencing, cell counts and neutrophil extracellular trap (NET) immunoassays were performed on bronchoscopic lower airway samples in 200 bronchiectasis subjects (108 NTM-, 92 NTM+). A preclinical model of oral commensal micro-aspiration and NTM infection was used to profile the murine lower airways with flow cytometry and a NET assay.

**Measurements and Main Results:** Lower airways of NTM+ bronchiectasis patients were enriched with *Mycobacterium* and oral commensals (e.g., *Veillonella, Prevotella*). NET levels were higher in NTM+ BAL. *Mycobacterium* and oral commensals co-occurred with NET and neutrophils in network studies. Distinct oral commensal taxa associated with severe disease phenotypes such as cavitary disease and exacerbators. In a murine micro-aspiration model, the combination of oral commensals and *Mycobacterium* led to a sustained pro-inflammatory immune response marked by an increase in Th17, γ8T cells, PD-1+ T lymphocytes as well as higher NET levels.

**Conclusions:** Our analyses showed that distinct microbiome features beyond the primary pathogen can contribute to neutrophilic inflammation and severe disease phenotypes in bronchiectasis/ NTM-LD.

## Introduction

Bronchiectasis is a heterogeneous and polymicrobial lung disease with a multitude of phenotypes (1). Majority of microbiome studies in bronchiectasis have used non-invasive sputum samples and therefore not profiled the lower airway micro-environment. Additionally, most studies have focused on cohorts that have a low prevalence of non-tuberculous mycobacterial lung disease (NTM-LD). In recent decades, NTM-LD has been on the rise globally (2), (3), translating into significant healthcare costs (4), and highlighting the need for rigorous pathogenesis studies.

Neutrophils extracellular traps (NET) are an emerging biomarker of lung inflammation. As web-like structures that can “trap” pathogens and prevent microbial dissemination (5), but excess NETosis can cause lung injury (6). In an international multi-cohort bronchiectasis study, higher NET levels associated with disease severity, lower alpha diversity and increased mortality, but the study did not include NTM+ subjects (6). Several other clinical trials in bronchiectasis have also excluded NTM+ patients (7–10). A recent study showed that *Mycobacterium avium* complex (MAC) can induce NETosis and NET-dependent production of matrix metalloproteinases, thus causing progressive inflammation(11). Here, we hypothesized that specific microbiome signatures such as enrichment with oral commensals can be associated with an aberrant neutrophil response and severe phenotypes in NTM+ bronchiectasis. Using bronchoscopic samples from 200 subjects (108 NTM-, 92 NTM+), we identified microbial (*Mycobacterium* and oral commensals) and host neutrophil signatures that associated with NTM positivity and severe clinical phenotypes. Then, using a murine model of NTM infection, we experimentally showed that exposure to both *Mycobacterium* and oral commensals triggered higher NET levels and a pro-inflammatory profile in both acute and chronic stages.

## Methods

Additional details on all methods are provided in the online data supplement.

### Study Design

This was a prospective observational study of 200 bronchiectasis patients enrolled at NYU Langone Medical Center (IRB# S14-01400). Inclusion criteria were imaging abnormalities and symptoms consistent with bronchiectasis. We excluded participants who had short-term antibiotics within 30 days of the bronchoscopy, subjects with pneumonia/active malignancy and subjects receiving immunoglobulins, suppressive macrolides, steroids or other biologics. In all subjects who underwent a bronchoscopy, bronchoalveolar lavage (BAL), supraglottic (Sup) and background/equipment (BKG) samples were collected.

### Clinical Data Indices

Disease severity was evaluated with the FACED Score (12), history of exacerbations in the previous year (13) (14), presence or absence of cavitary disease and a detailed Chest CT score. Diagnosis of NTM-LD was made with the American Thoracic Society (ATS) guidelines (15).

### Lower airway taxonomy using 16S rRNA gene sequencing

Microbiome diversity indices were evaluated by alpha and beta diversity measures. Differential taxonomic enrichment was analyzed with EdgeR package (v3.36.0) (16–18). Microbiome ‘metacommunities’ were determined with Dirichlet Multinomial Mixture Modeling (19). Microbe-microbe, microbe-cell and microbe-NET associations were evaluated with co-occurrence analyses (20) (21) and MaAsLin2 (22).

### Aerosol NTM infection in a BALB/c microaspiration model

Micro-aspiration was performed with mixed oral commensals (referred to as MOC) composed of *Streptococcus mitis, Veillonella parvula* and *Prevotella melaninogenica* (23, 24). Aerosol NTM infection was performed with MAC101 strain. Experimental arms are illustrated in Fig. 6A. Readouts included flow cytometry and BAL fluid (BALF) NET ELISA at 1 and 2 months after NTM infection.

#### NET ELISA

Human and murine BALF NET ELISA have been described in the online data supplement (25), (26).

### Statistical analyses

Mann-Whitney test and Kruskal-Wallis ANOVA (in case of > 2 categories) were used for non-parametric tests of association. Wilcoxon signed-rank test was used for paired analyses.

## Results

### Demographics, disease severity indices, radiologic phenotypes and BALF NET levels

In table 1, we summarize demographics, disease severity indices, radiologic phenotypes and NET levels. Average age of the cohort was 67, with a female predominance (67% in NTM-versus 81% in NTM+, p=0.02, Mann-Whitney). A multivariable logistic regression analysis with covariate adjustment for age, sex, race and body mass index (BMI) showed that lower BMI (OR =0.92; 95% CI: 0.85-0.99; p = 0.028) and older age (OR =1.02; 95% CI: 1.00-1.05; p = 0.049) were independently associated with increased odds of NTM positivity. Ninety-two subjects (46%) met the ATS NTM-LD diagnostic criteria (15). Approximately 33% of all subjects had a history of gastroesophageal reflux disease, regardless of NTM status. No statistically significant differences were noted for other co-morbidities between the two groups. Respiratory physiological parameters were also similar with an obstructive component in 37.5% of the cohort.

**Table 1:**
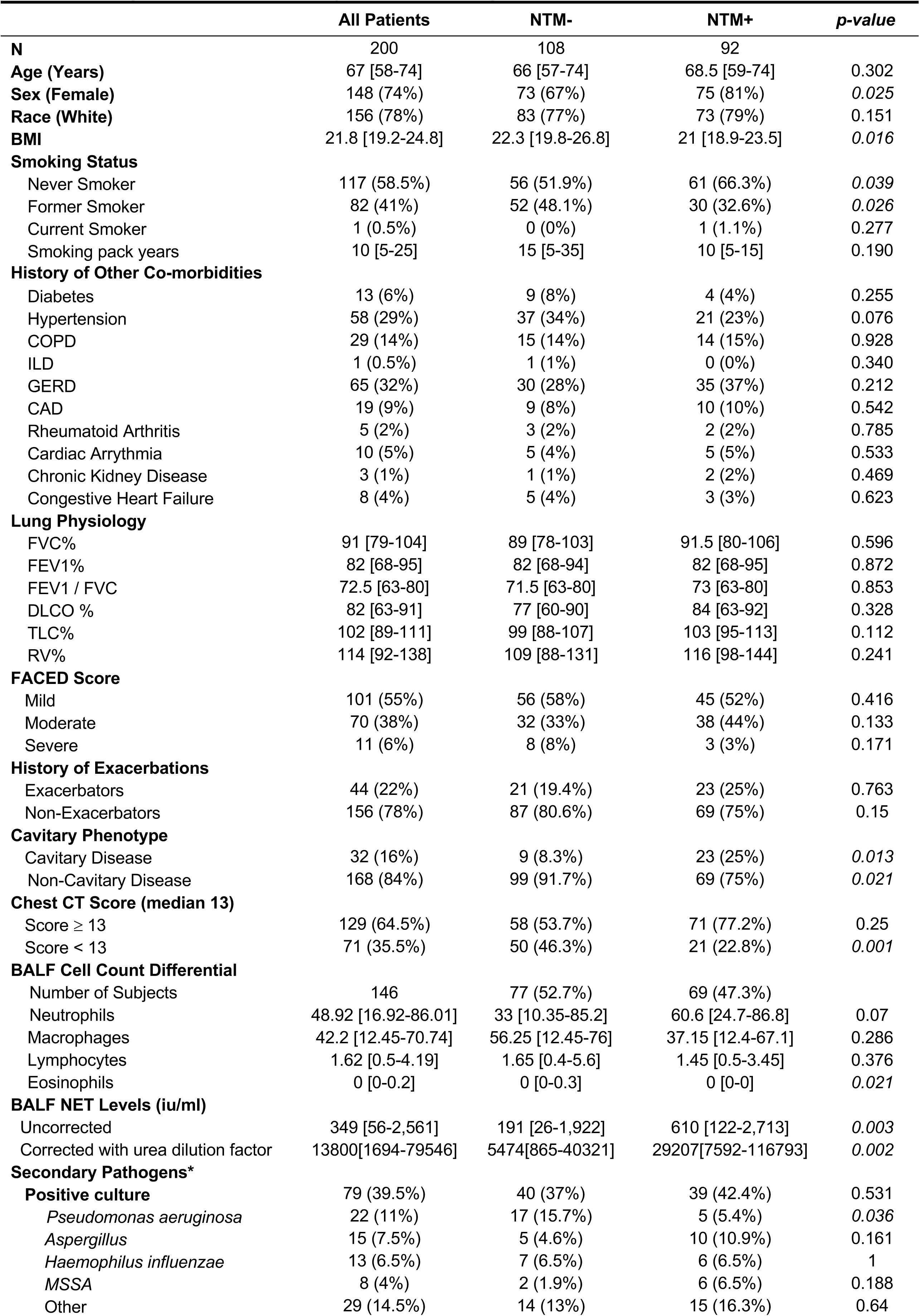
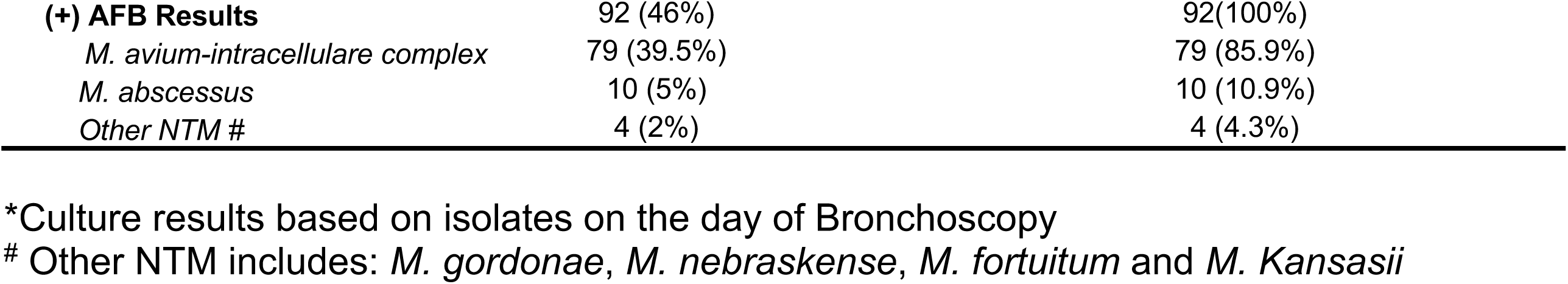
Demographics, disease severity indices, radiologic phenotypes and BALF NET levels. In a cohort of 200 subjects with bronchiectasis, 92 (46%) met the ATS NTM-LD diagnostic criteria. Average age was 67 with a female predominance (67% in NTM-versus 81% in NTM+, p=0.02, Mann-Whitney). Approximately 33% of all subjects had a history of gastroesophageal reflux disease. No statistically significant differences were noted for other co-morbidities between the two groups. Respiratory physiological parameters were similar between the groups with an obstructive component in 37.5% of the cohort. There was no significant difference in FACED scores and history of exacerbations between the NTM-/NTM+ groups. Radiologically, 23/92 (25%) of NTM+ subjects had a cavitary phenotype, as compared to 9/108 (8.3%) of NTM-subjects (p=0.013, Mann-Whitney). Median Chest CT score was 13 [IQR: 12-16], with a higher proportion of NTM+ subjects who had a higher-than-median Chest CT score but this did not reach statistical significance. BALF NET levels were at a median value of 349 iu/ml [IQR 56-2561], with significantly higher levels in the NTM+ group compared to the NTM-group (p=0.003, Mann-Whitney, Table 1). After standardization with the urea dilution method, the NET levels had a median value of 13800 iu/ml [IQR 1694-79546], still with significantly higher levels in the NTM+ group (p=0.002, Mann-Whitney, Table 1). BALF cell count differentials were similar in both groups. A secondary respiratory pathogen was isolated on the day of the bronchoscopy in 37% of subjects in the NTM-group, and in 42% in the NTM+ group. The commonest organisms cultured were *Pseudomonas aeruginosa, Aspergillus spp.* and *Haemophilus influenzae*, with *Pseudomonas* being the most abundant secondary pathogen in the NTM-group. Majority of the mycobacterial isolates were identified as MAC complex (80/92 or 85%).

More than half the subjects (55%) had a mild FACED score, followed by moderate (38%) and severe (6%) scores. There was no significant difference in FACED scores and history of exacerbations between the NTM-/NTM+ groups. Radiologically, 23/92 (25%) of NTM+ subjects had a cavitary phenotype, as compared to 9/108 (8.3%) of NTM-subjects (p=0.013, Mann-Whitney). Median Chest CT score was 13 [IQR: 12-16], median cavity size was 2.4 cm [IQR 2-4] and median number of cavities was 1 [IQR 1-4.25]. A higher proportion of NTM+ subjects had a higher-than-median CT score (77.2% as compared to 53.7% of NTM-subjects) but this did not reach significance. BALF NET levels had a median value of 349 iu/ml [IQR 56-2561], with significantly higher levels in the NTM+ group compared to the NTM-group (p=0.003, Mann-Whitney). After standardization with the blood/BAL urea dilution factor, NETs had a median value of 13800 iu/ml [IQR 1694-79546], and with significantly higher levels in the NTM+ group (p=0.002, Mann-Whitney). BALF cell count differentials were similar in both groups.

### Microbiota comparison across airways using bronchoscopic and supraglottic samples

16S rRNA gene sequencing was performed on BAL samples from the radiologically most diseased lung segment in every subject. A subset of supraglottic (Sup, n=170) and background/control (BKG, n=36) samples were included. Bacterial load as measured by droplet digital PCR (ddPCR), was lowest in BKG samples, followed by BAL samples, and was highest in the Sup samples (p<0.001 for all comparisons, Mann-Whitney, Fig. E1A). Total reads were also lowest in BKG samples, followed by BAL samples and highest in Sup samples (BKG versus Sup, p<0.001, Mann-Whitney, Fig. E1B). The median sequence depth available for downstream analyses was 24,829 [IQR=12,872-51,844] reads per sample. Spearman’s rank correlation coefficient between BAL bacterial load and NET levels was highly significant, regardless of NTM status (Spearman’s p= 5.08e-15, Fig. E1C). Alpha diversity (Shannon index) was lower in the Sup samples than in BAL samples (p=0.038, Mann-Whitney, Fig. E1D). Beta diversity analyses (Bray Curtis) showed clear compositional differences between the three sample types (p=0.001, PERMANOVA, Fig. E1E). Due to low biomass of BAL samples, we evaluated potential bacterial DNA contaminants in our dataset using Decontam (Figure E2A-C, and Table E1). Given the compositional nature of the data, and challenges in confirming the “contaminant” nature of taxa, we did not remove these taxa from subsequent analyses, rather labelled them as potential contaminants for reference. Finally, Fig. E3 shows that BAL had a higher relative abundance of *Haemophilus, Ralstonia, Flavobacterium* and *Pseudomonas*, compared to Sup samples.

### Comparison of lower airway microbiota in NTM-/NTM+ subjects

There were no significant differences in the bacterial burden, alpha diversity or beta diversity between NTM-/NTM+ subjects (Fig. 1A-C). Taxonomically, NTM-BAL had a higher relative abundance of *Moraxella, Enterobacteriaceae, Streptococcus* and *Actinobacillus,* whereas NTM+ BAL had a higher relative abundance of *Proteus, Stenotrophomonas, Actinomycetes* and *Nocardia* (Fig. 1D). In our cohort, 24/92 (26%) of NTM+ subjects were either on NTM treatment at the time of bronchoscopy or were treated in the past. To eliminate confounding effects of antibiotics on the microbiome analyses, we undertook a subgroup analysis with the 68/92 (74%) NTM+ subjects who had never been treated with any first/second/third line anti-NTM agent. Similar to the full cohort, we found no significant differences in the bacterial burden, alpha diversity or beta diversity between NTM-/NTM+ never-treated subjects (Fig. E4A-C). Taxonomically, never-treated NTM-BAL had a higher relative abundance of *Moraxella, Enterobacteriaceae, Streptococcus* and *Actinobacillus,* and never-treated NTM+ BAL had a higher relative abundance of *Proteus, Stenotrophomonas, Actinomycetes* and *Nocardia* (Fig. E4D).

**Figure 1.**
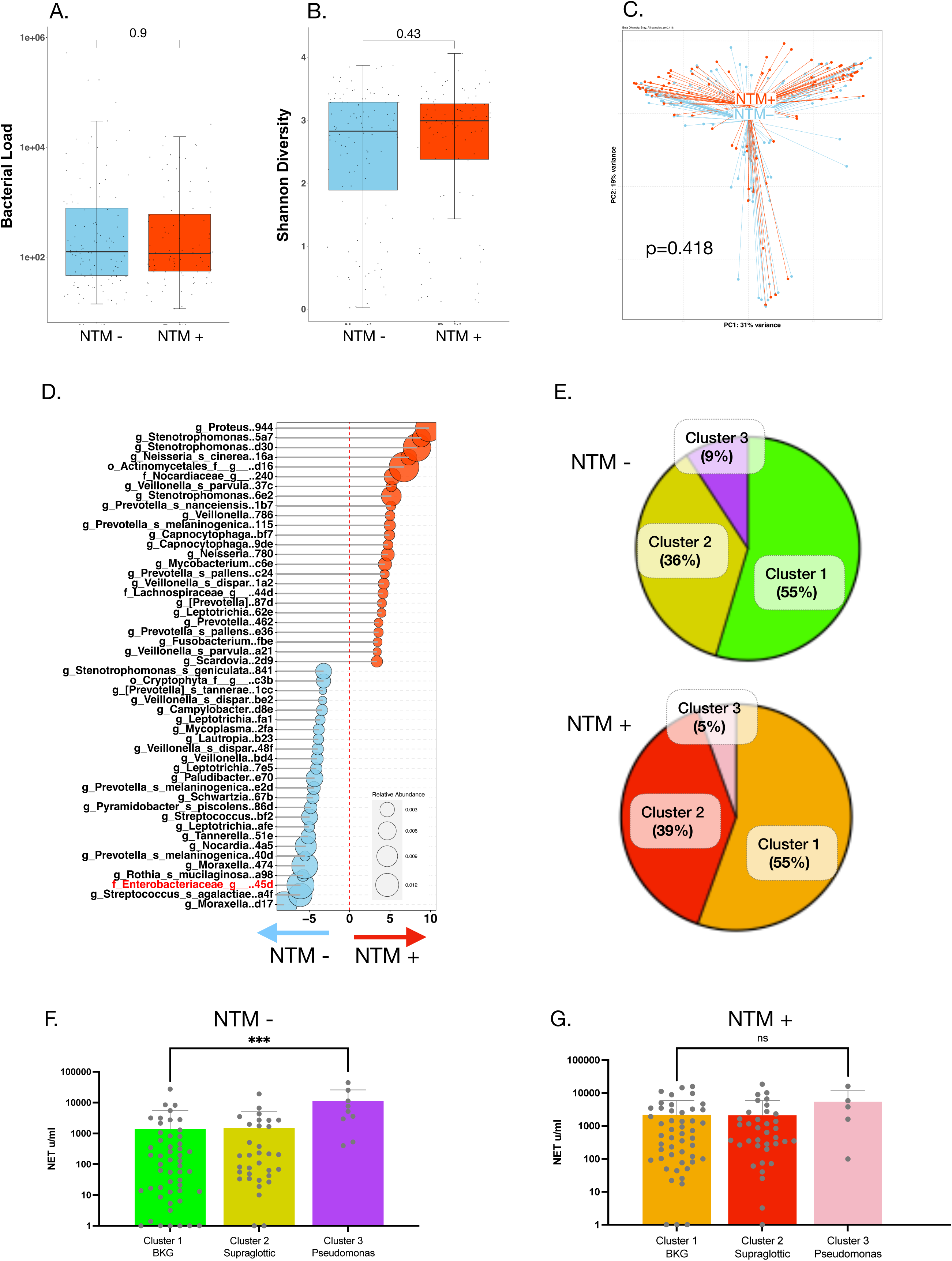
Comparison of the lower airway microbiota in NTM-/ NTM+ subjects. Fig. 1A**-C**. There were no significant differences in the bacterial burden, alpha diversity (Shannon) or beta diversity (Bray Curtis Dissimilarity Index) between NTM-/NTM+ subjects. Fig. 1D. Taxonomically, NTM-BAL samples had a higher relative abundance of *Moraxella, Enterobacteriaceae, Streptococcus* and *Actinobacillus*, whereas NTM+ BAL had a higher relative abundance of *Proteus, Stenotrophomonas, Actinomycetes* and *Nocardia.* Fig. 1E. Dirichlet Multinomial Modeling identified that more than 50% of the lower airway samples belonged to cluster 1, <40% to cluster 2 and <10% to cluster 3, with relatively similar proportions in both NTM-/NTM+ groups. Fig. 1F**-G**. Lower airway NET levels were significantly higher in cluster 3 in NTM-subjects (p<0.001, Mann-Whitney) but there were no statistically significant differences in NET levels in the NTM+ clusters.

### Taxonomic overlap of cultured pathogens and 16S rRNA gene sequencing

Within the NTM+ group, majority of the mycobacterial isolates were MAC complex (79/92 or 39.5%), with a small number of *Mycobacterium abscessus, Mycobacterium fortuitum* and *Mycobacterium kansasii* (Table 1). Consistent with our prior report (27), only 38% of NTM+ subjects had any 16S rRNA sequence reads annotated to *Mycobacterium* (35/92) (Fig. 2A). The majority showed low relative abundance (<10%), with most abundant ASVs identifying *Mycobacterium avium* as the top potential strain. In the NTM culture negative group, five subjects had mycobacterial 16S rRNA reads, but the relative abundances were extremely low (<1%, Fig. 2B). Blast analyses for four subjects were consistent with a non-pathogenic *Mycobacteria* (Table E2). One NTM-subject who had a high relative abundance (>30%) of *Mycobacterium avium* had been on protracted antibiotics, with a negative BAL culture immediately before surgery, and therefore the NTM-label. Secondary respiratory pathogens were isolated on the day of bronchoscopy in 37% of NTM-subjects, and in 42% of NTM+ subjects (Table 1). The commonest organisms cultured were *Pseudomonas aeruginosa, Aspergillus spp.* and *Haemophilus influenzae*. In the NTM-group, *Pseudomonas aeruginosa* was the most abundant secondary pathogen with a statistically higher prevalence than in NTM-group (15.7% vs 5.4%, Chi-square, p=0.036, Table 1). Abundance of amplicon sequence variants (ASV)s annotated to *Pseudomonas* and *Haemophilus* had a strong concordance with culture positivity (Fig. E5A-B), but this was less clear for *Streptococcus* (Fig. E5C). Receiver-operator characteristic curves comparing 16S rRNA with culture positivity showed an AUC of 0.95 and 0.98, respectively, for *Pseudomonas* and *Haemophilus* (Fig. E6A-B) and an AUC of 0.54 for *Streptococcus* (Fig. E6C). Direct comparison of the relative abundances of *Pseudomonas, Haemophilus* and *Streptococcus* with NET levels showed a highly significant Spearman’s rank correlation coefficient for *Pseudomonas* (Spearman’s p=8.96e-05, Fig. E6D), but it was not significant for *Haemophilus* or *Streptococcus* (Fig. E6E-F).

**Figure 2.**
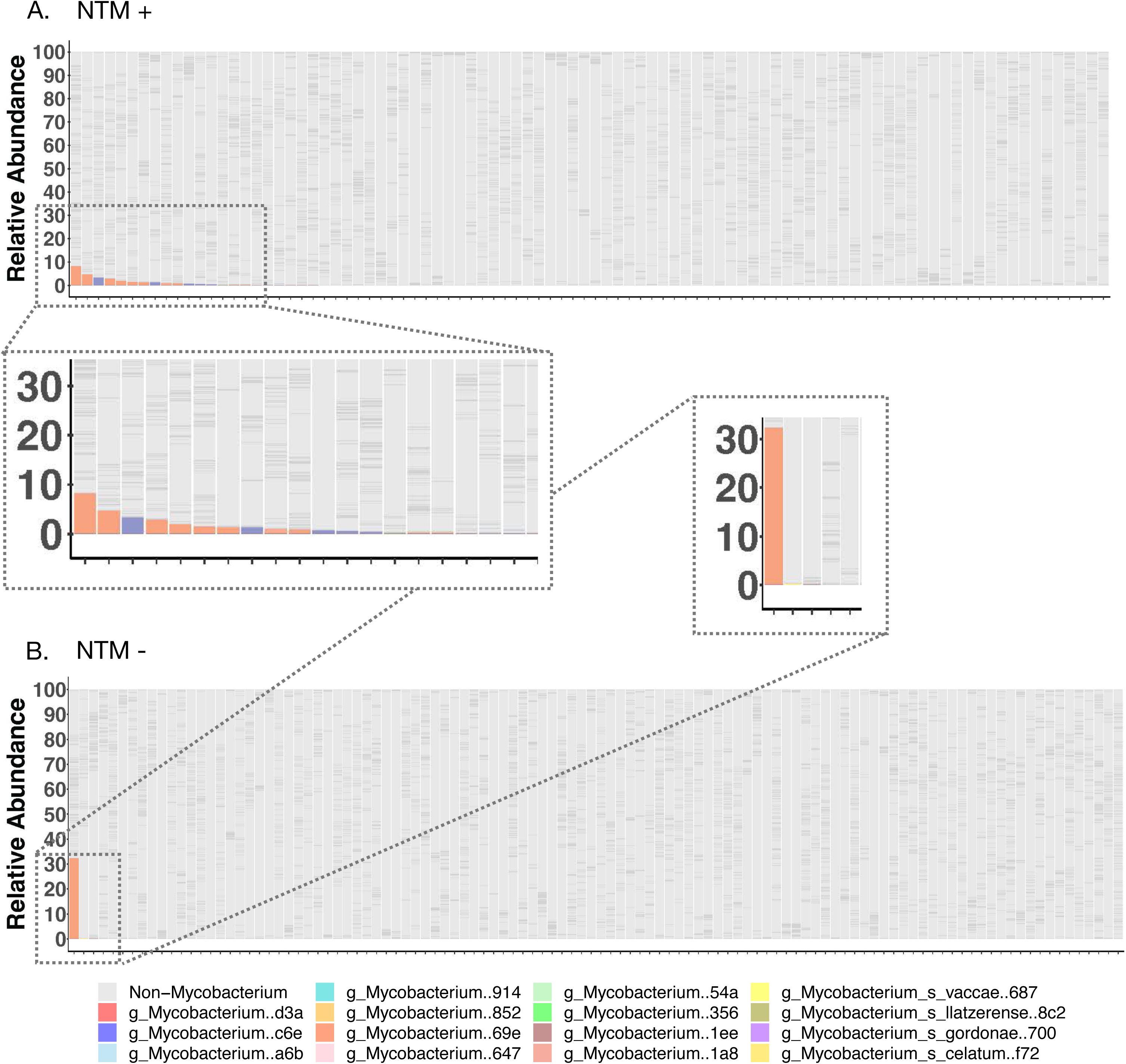
Taxonomic overlap of cultured pathogens and 16S rRNA gene sequencing. Fig. 2A. In the NTM+ group, 38% of subjects had sequence reads annotated to *Mycobacterium* (35/92) with the vast majority showing low relative abundances (<10%). The most abundant ASVs identified *Mycobacterium avium* as the top potential strain. Fig. 2B. In the NTM-group, BAL samples from five subjects had some mycobacterial reads in their 16S rRNA gene sequencing data, but the relative abundances were extremely low (<1%).

### Lower airway metacommunities in NTM-/NTM+ subjects

Using DMM clustering, we identified three possible clusters that had the best fit for the data (Fig. E7). Bacterial load was significantly different between all three clusters (1 versus 2: p<0.0001; 2 versus 3: p=0.00031; 1 versus 3: p<0.0001, Mann-Whitney, Fig. E8A). Cluster 1 had the lowest bacterial load whereas cluster 3 had the highest. Alpha diversity was also significantly different between the clusters (2 versus 3: p<0.0001 and 1 versus 3: p<0.0001, Mann-Whitney, Fig. E8B). Cluster 1 and 2 had higher alpha diversities than cluster 3. Bray Curtis analyses showed all three clusters to be compositionally distinct (p<0.0001, PERMANOVA, Fig. E8C). Cluster 1 had a high relative abundance of background taxa, cluster 2 was enriched with oral commensals (Fig. E8D-E) and cluster 3 was enriched with dominant pathogens (Fig. E8F). More than 50% of the lower airway samples belonged to cluster 1, <40% to cluster 2 and <10% to cluster 3, with relatively similar proportions in both NTM-/NTM+ groups (Fig. 1E). Categorization of 16S rRNA taxonomic annotation based on pathogenic potential at the genus level (Fig. E9) showed significant overlap with DMM clusters, with 32% of the “pathogens (P)” overlapping with Cluster 3 (*Pseudomonas-*dominant), 85% of the “possible-pathogens (PP)” overlapping with cluster 2 (supraglottic-dominant), and 92% of “non-pathogens (NP)” overlapping with cluster 1 (background), (Chi-Square p< 2.2e-16). Differences in alpha diversity were significant between the NP, PP and P groups; with the lowest value for the P and highest for the NP cluster (p<0.0001 for all comparisons, Mann-Whitney, Fig. E9A). Beta diversity was also significantly different between the three clusters (p<0.0001, PERMANOVA). The NP group was enriched with *Flavobacterium* (Fig. E9C), the PP group was enriched with *Prevotella* and *Veillonella* (Fig. E9D), and the P group (lowest alpha diversity) was enriched with *Pseudomonas*, *Haemophilus* and *Streptococcus* (Fig. E9E). Table E3 shows the classification of all taxa into pathogenic, possible pathogenic or non-pathogenic species.

### Oral commensals associate with NET in NTM+ lower airways

Lower airway NETs were significantly higher in NTM-cluster 3 (Fig. 1F, p<0.001) but there were no statistically significant differences in the NTM+ clusters (Fig. 1G). Cluster 3 was associated with a higher neutrophil and lower macrophage cell count as compared to cluster 1 and cluster 2 (Fig. E10A-B). A biplot analysis of the distribution of taxa in relation to inflammatory cells and NETs showed that *Pseudomonas* was associated with higher NETs and neutrophils in the NTM-group (Fig. 3A), whereas such an association was noted for *Mycobacterium* and the oral commensals *Megasphera and Neisseria* in the NTM+ group (Fig. 3B). In microbial-cellular interaction networks (20), *Pseudomonas* competed against oral commensals (negative correlation; red lines) such as *Fusobacterium, Rothia, Actinomyces* and *Campylobacter;* but had a synchronous and interdependent relationship with neutrophils (green line, Fig. 3C) and a strong association with NETs (MaAsLin2 coefficient of association, CoA, 0.67) (Fig. 3E). In contrast, in NTM+ subjects, *Mycobacterium* had a synchronous interaction with oral commensals such as *Chryseobacterium, Capnocytophaga, Fluviicola* (green lines, Fig. 3D) and associated with NETs (MaAsLin2, CoA 0.18) (Fig. 3F).

**Figure 3.**
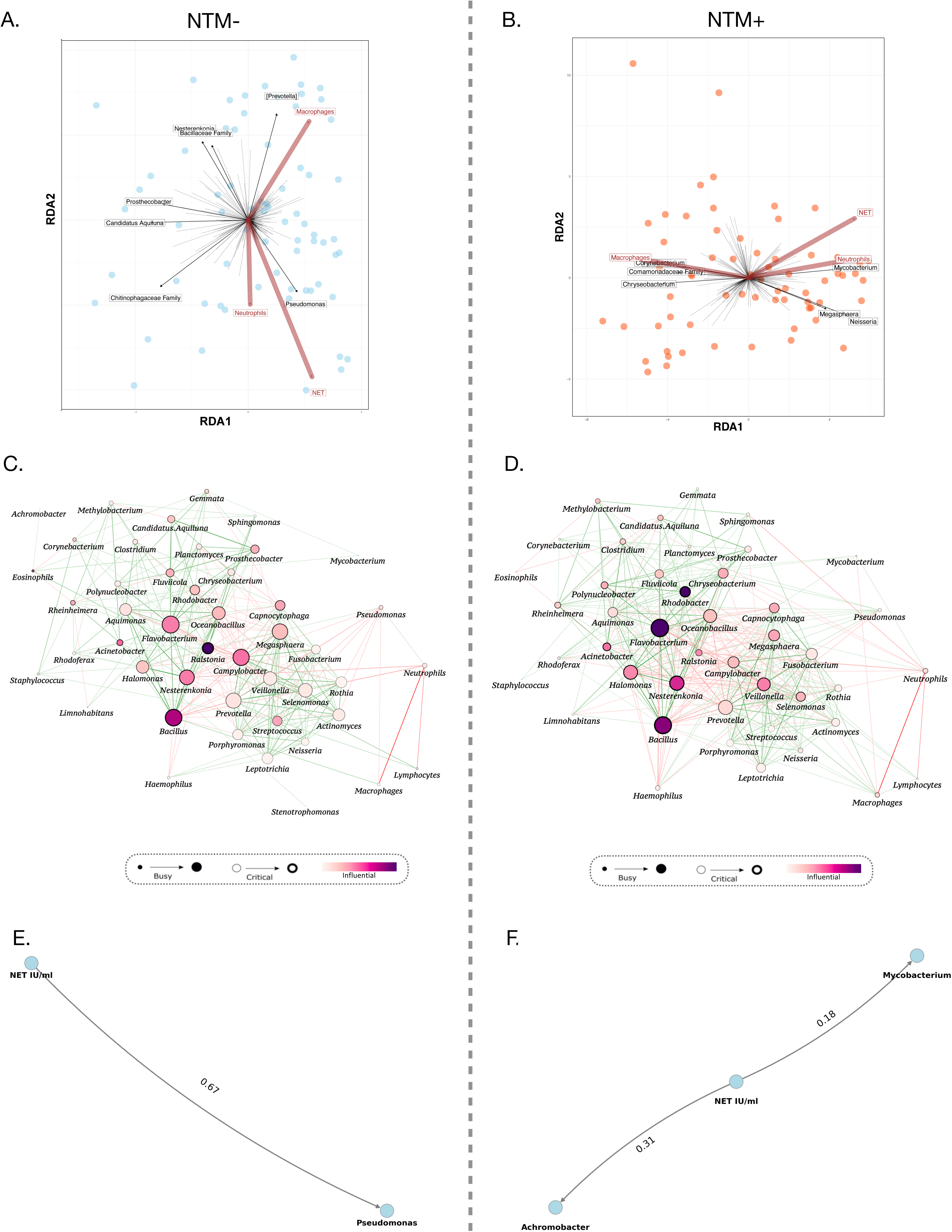
*Pseudomonas* and *Mycobacteria* co-occur with neutrophils and NET. Fig. 3A. On a biplot analysis, *Pseudomonas* was associated with higher NET levels and neutrophils in NTM-BAL. Fig. 3B. In the NTM+ group, *Mycobacterium* and oral commensals were associated with NET and neutrophils. Fig. 3C. In microbial-cellular interaction networks, *Pseudomonas* in NTM-subjects competed against oral commensals (negative correlation; red lines). Fig. 3D. In NTM+ subjects, *Mycobacterium* had a synchronous interaction with oral commensals (positive correlation, green lines). Fig. 3E**-F**. MaAsLin2 analysis showed a strong association of *Pseudomonas* and *Mycobacteria* with NETs (coefficients of association, 0.67 and 0.18, respectively).

### Oral commensal signatures dominate in NTM+ severe phenotypes

We explored microbial signatures in severe disease phenotypes based on Chest CT score, cavitation and exacerbator profiles. Due to the small number of patients in the severe FACED category (Table 1), we did not use this score for our analyses. Using Chest CT scores, we found compositional differences in NTM-/NTM+ subjects (p=0.017 and 0.012 respectively, PERMANOVA, Fig. E11A-B), and a higher relative abundance of *Haemophilus, Achromobacter* and oral commensals such as *Veillonella, Prevotella, Stenotrophomonas*, *Proteus* in the NTM+ “above-median-CT-score” group (Fig. E11D). The NTM-“above-median-CT-score” group showed some *Nocardia* and *Streptococcus* ASVs (Fig. E11C). We then compared cavitary versus non-cavitary subjects and identified significant compositional differences in both NTM-/NTM+ groups (p=0.009 and 0.013 respectively, PERMANOVA, Fig. 4A-B). Taxonomically, the NTM-cavitary microbiome had a higher relative abundance of *Pseudomonas* (Fig. 4C), whereas the NTM+ cavitary microbiome showed ASVs from *Haemophilus* and *Streptococcus* (Fig. 4D). Finally, we found unique signatures in the exacerbator/non-exacerbator sub-cohort. Beta diversity between exacerbators/non-exacerbators were divergent in both NTM-/NTM+ groups (p=0.021 and 0.011 respectively, PERMANOVA, Fig. 5A-B). There was a preponderance of *Moraxella* and *Prevotella* in NTM-exacerbators (Fig. 5C), and a preponderance of *Streptococcus* and *Achromobacter* in NTM+ exacerbators (Fig. 5D).

**Figure 4.**
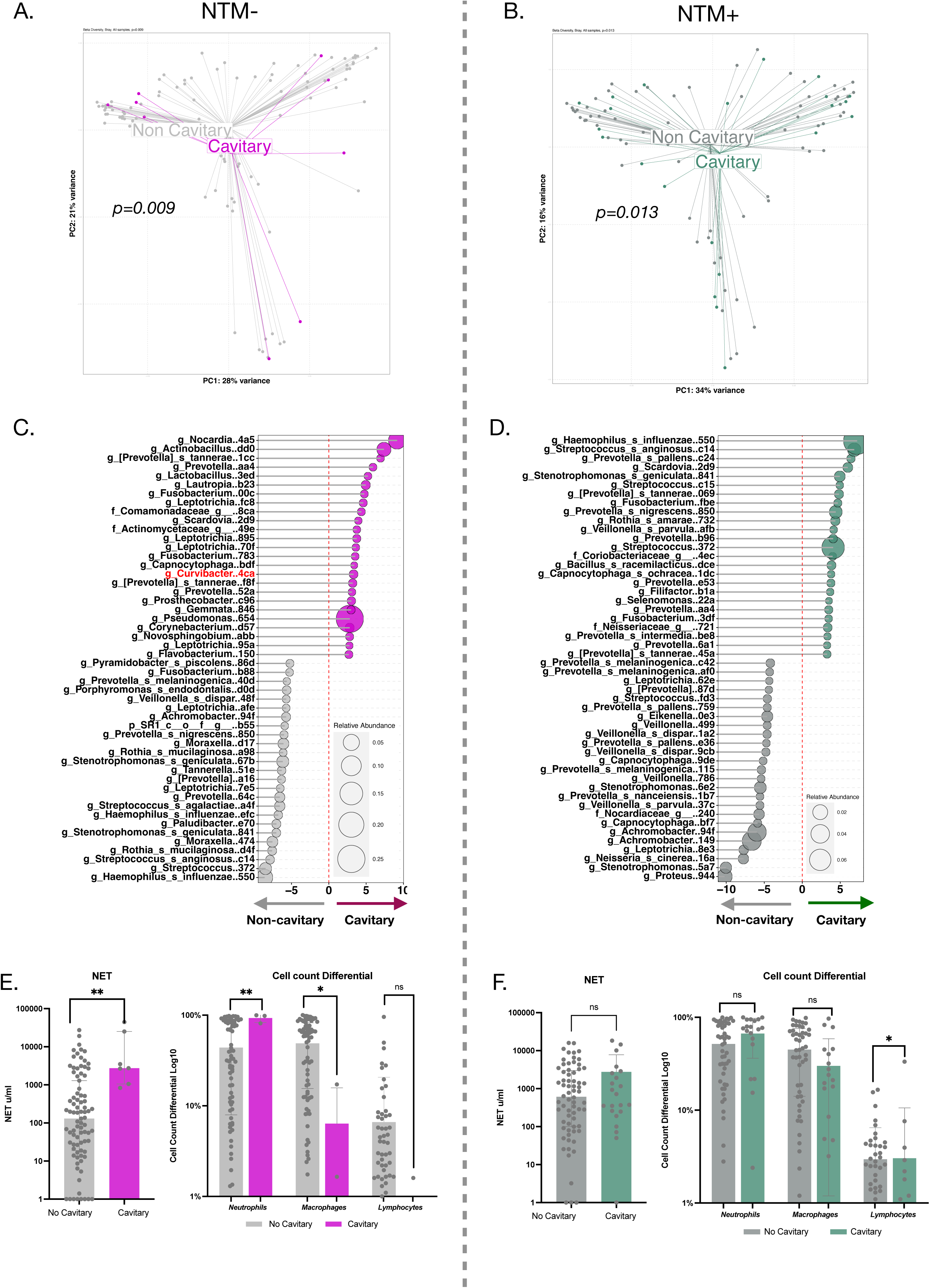
Microbial signatures associate with disease severity (cavitary profile). Fig. 4A, B. Beta diversity plots showed that cavitary disease subjects had significant compositional differences from those without cavitary disease, in both NTM-/NTM+ groups (p=0.009 and 0.013 respectively, PERMANOVA). Fig. 4C. Taxonomically, the NTM-cavitary microbiome had a higher relative abundance of *Pseudomonas* whereas NTM+ cavitary microbiome showed ASVs from *Haemophilus* and *Streptococcus* (Fig. 4D). NET levels were significantly higher in NTM-patients with a cavitary phenotype (Fig. 4E) but this difference was not statistically significant in the NTM+ group (Fig. 4F). A statistically significant difference was also noted for neutrophil and macrophage cell count differentials in the NTM-group (Fig. 4E, F).

**Figure 5.**
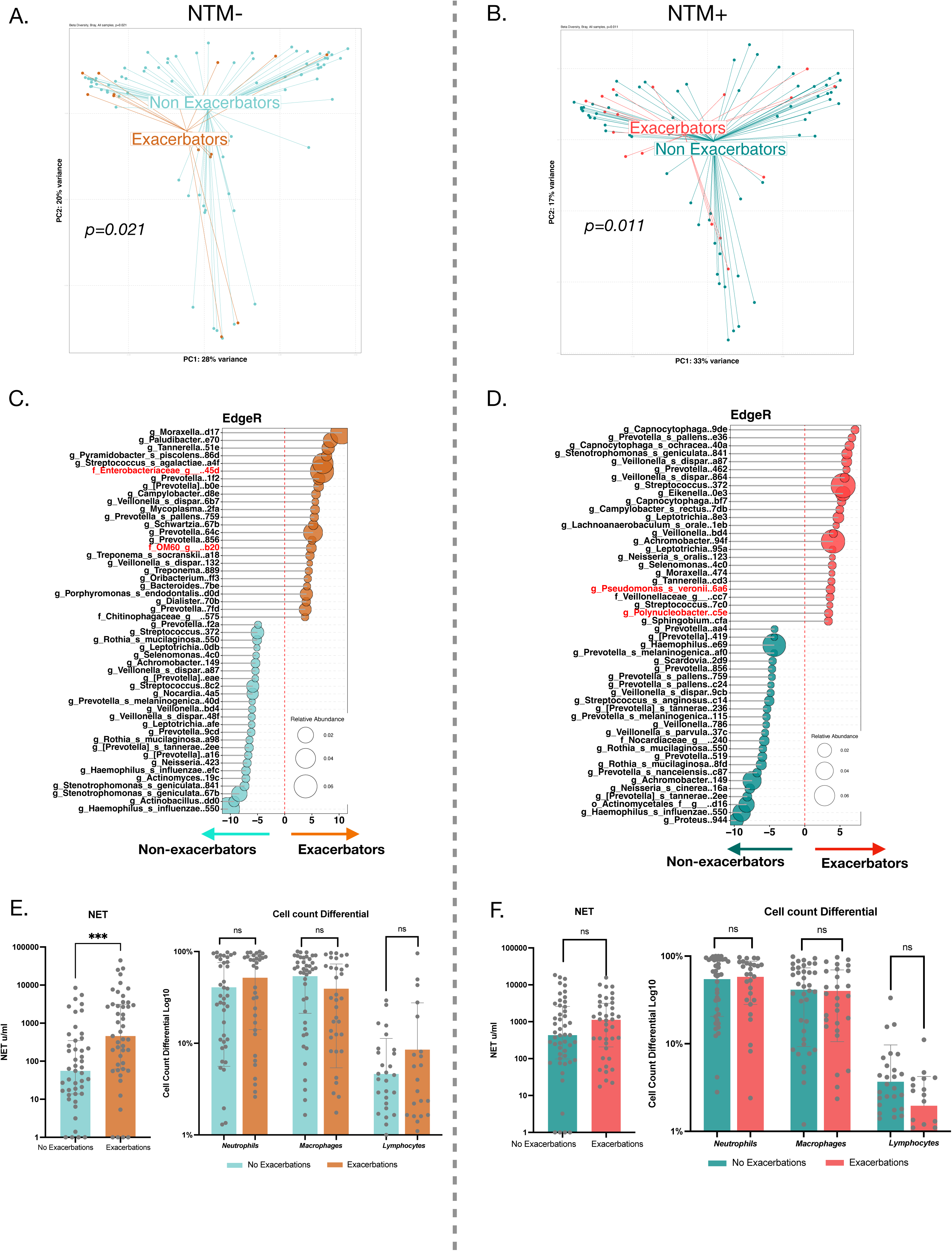
Microbial signatures associate with disease severity (exacerbator profile). Fig. 5A, B. Beta diversity between exacerbators/non-exacerbators were divergent in both NTM-/NTM+ groups (p=0.021 and 0.011 respectively, PERMANOVA). There was a preponderance of *Moraxella* and *Prevotella* in NTM-exacerbators (Fig. 5C), and a preponderance of *Streptococcus* and *Achromobacter* in NTM+ exacerbators (Fig. 5D). NET Levels were significantly higher in the exacerbator phenotype within the NTM-group (Fig. 5E) but this difference was not statistically significant in the NTM+ group (Fig. 5F). Cell count differentials did not show any statistically differences in either the NTM- or in the NTM+ groups (Fig. 5E**-F**).

### NETs co-occur with oral commensals in severe phenotypes

In the cavitary microbiome, NETs had a positive interaction with *Pseudomonas* (CoA of 0.93, MaAsLin2, Fig. E12A). *Pseudomonas* exhibited competitive interactions with oral commensals (red lines) whereas *Mycobacteria* showed synchronous interactions with oral commensals (green lines) (Fig. E12B). NETs were significantly higher in NTM-cavitary phenotype (Fig. 4E) and significant differences were noted for neutrophil and macrophage cell count differentials in the NTM-cavitary group, but not in the NTM+ group (Fig. 4E-F). In exacerbators, NETs had a positive interaction with *Pseudomonas* and *Achromobacter* (MaAsLin2, CoA 0.62 and 0.24, respectively) and a negative interaction with *Flavobacterium* (MaAsLin2, CoA -0.42) (Fig. E12C). Similar to the cavitary microbiome, *Pseudomonas* had a competitive interaction (red lines) whereas *Mycobacteria* had a synchronous interaction (green lines) with oral commensals in exacerbators (Fig. E12D). NETs were significantly higher in NTM-exacerbators (Fig. 5E). Cell count differentials did not show any change (Fig. 5E-F).

### NTM drives a NET and Th-17/neutrophilic response in a murine micro-aspiration model

Having shown the associations between *Mycobacterium*, oral commensals and NETs in the human cohort, we investigated the effects of micro-aspiration with MOC in a murine NTM infection model. Significantly increased murine BALF NET levels were noted both at 1 and 2 months after NTM infection, with highest levels in the NTM+MOC group (p<0.0001 at 1 and 2 months, One-way ANOVA, Fig. 6B). Importantly, NETosis was significantly higher in the NTM+MOC group than the NTM-only-group at 2 months (p<0.0001, One-way ANOVA, Fig. 6B), suggesting that chronic exposure to oral commensals augmented neutrophil dysfunction. On FACS analyses, neutrophils were elevated in the NTM+MOC group at 1 month (p<0.0001, One-way ANOVA, Fig. 6C). Neutrophils down-trended thereafter, but still remained high at 2 months. Th-17 cells are dysfunctional in NTM-LD(28–30), and are associated with NETosis(31–33). Both Th-17 and γ8T cells drive sustained neutrophil activation(34, 35). Th-17 cells were significantly higher in the NTM+MOC group compared to the NTM-only-group at both 1 and 2 months (p=0.0231 and p<0.0001, respectively, One-way ANOVA, Fig. 6D). A similar trend was noted for γ8T cells in the NTM+MOC group at both time points (p=0.0002 and p<0.0001, respectively, One-way ANOVA, Fig. 6E). NTM infection also drove the influx of dendritic cells and macrophages at 1 month (p<0.0001 and p=0.0012, respectively, One-way ANOVA, Fig. E13A-B) and 2 months (p<0.0001 and p=0.0003, respectively, One-way ANOVA, Fig. E13A-B). This NTM-driven influx was augmented by MOC at 1 month for dendritic cells (p<0.0001, One-way ANOVA, Fig. E13A) and both at 1 and 2 months for macrophages (p=0.0022 and p=0.0047, respectively, One-way ANOVA, Fig. E13B). No significant trends were noted at either time points for eosinophils, monocytes, CD4+ T cells or Th1 cells (Fig. E13C-F). Both T_REG_ cells and CD8+ T cells seemed to down-trend, but this was not significant (Fig. E13G-H).

**Figure 6.**
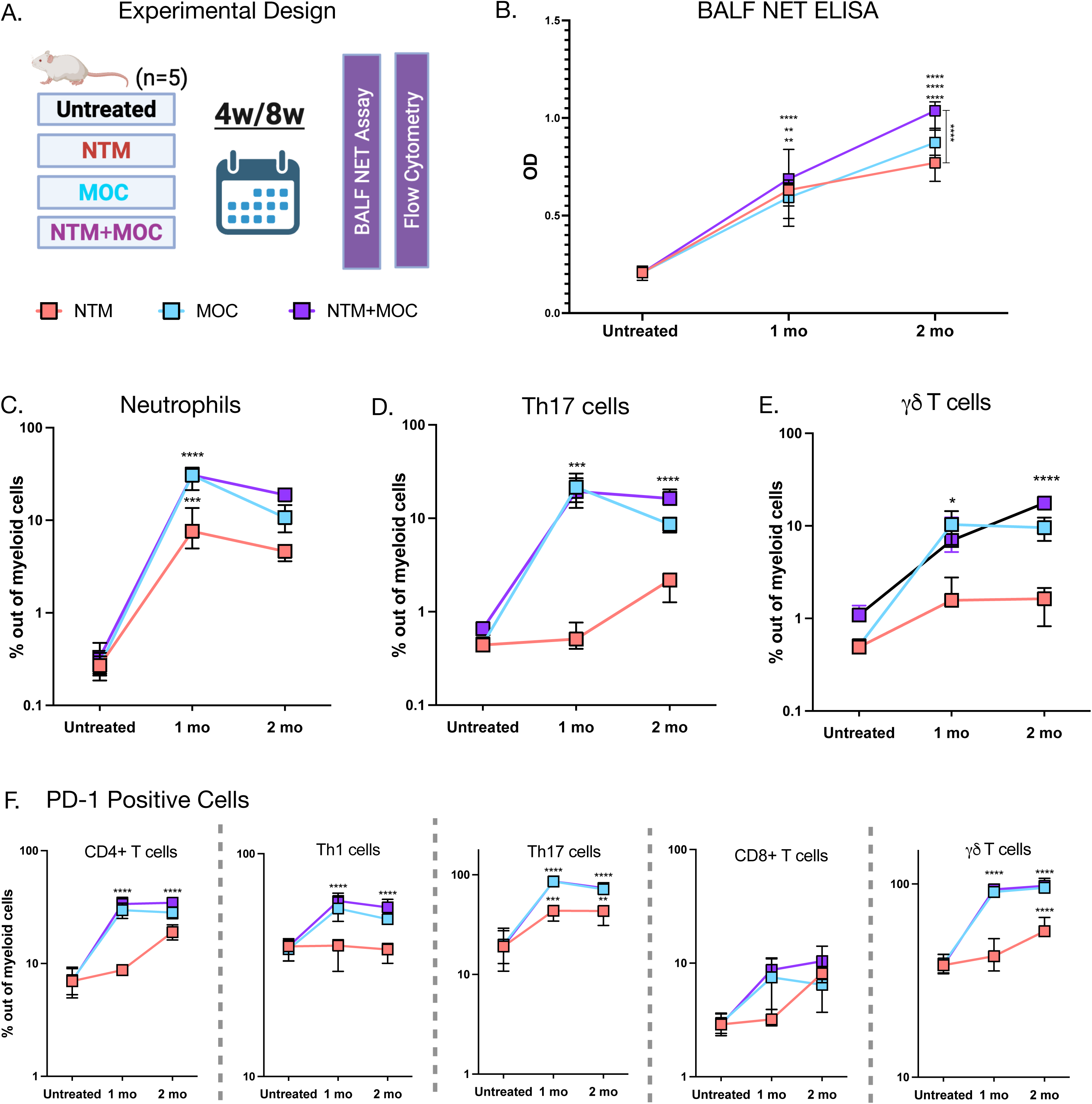
NTM drives a NET and Th-17/neutrophilic response in a murine micro-aspiration model. Fig. 6A. The four experimental groups were A: Untreated/control, B: Aerosol NTM infection only, C: Intratracheal Weekly MOC, D: NTM infection and weekly MOC. Mice were euthanized at 1 month or 2 months after NTM infection. Fig. 6B. Increased murine BALF NET levels were noted at 1 and 2 months after NTM infection, with the highest levels in the NTM+MOC group (p<0.0001 at both 1 and 2 months, One-way ANOVA). NETosis was further augmented in the NTM+MOC group compared to the NTM only group at 2 months (p<0.0001, One-way ANOVA). Fig. 6C. Neutrophils (CD11b+ Ly6G+) were significantly elevated in the NTM+MOC group at 1 month (p<0.0001, One-way ANOVA) and although neutrophils down-trended thereafter, they remained high at 2 months. Fig. 6D. Th-17 cells (CD4+ IL17A+ RORγt+) were significantly higher in the NTM+MOC groups when compared to the NTM group at both 1 and 2 months (p=0.0231 and p<0.0001, respectively, One-way ANOVA). Fig. 6E. An increase was also noted for γ8T cells (CD3+ TCRγ8+) in the NTM+MOC group at both time points (p=0.0002 and p<0.0001, respectively, One-way ANOVA). Fig. 6F. Significantly increased expression of PD-1 was noted in CD4+, Th1, Th17 and γ8T cells in the NTM+MOC group at both time points (p<0.0001 for all 4 cell types, One-way ANOVA). In contrast, NTM increased PD-1 expression on Th17 and γ8T cells only (p<0.0001 for both cell types at 2 months, One-way ANOVA).

Expression of PD-1 receptors is associated with smear-positive and radiologically severe NTM infection (36). We found significantly increased expression of PD-1 in CD4+, Th1, Th17 and γ8T cells in the NTM+MOC group at both time points (p<0.0001 for all 4 cell types at both time points, One-way ANOVA, Fig. 6F). In contrast, the NTM-only-group had increased PD-1 expression on Th17 and γ8T cells only (p<0.0001 for both cell types at 2 months, One-way ANOVA, Fig. 6F). In summary, our mouse data supports that NTM infection and chronic oral commensal exposure can drive a sustained inflammatory response and high NET levels.

## Discussion

NTM lung infection is a difficult to treat condition that involves a complex interplay between microbial and host factors (15). Protracted antibiotic regimens have translated into high healthcare costs and highlight the pressing need for adjunctive, host-directed interventions. In this study, using bronchoscopic samples from a cohort of 200 bronchiectasis subjects we showed that a lower airway microbiome enriched with oral commensals was associated with NTM positivity and clinical disease severity. While the dominance of *Proteobacteria* has long been associated with poor outcomes and mortality in non-NTM bronchiectasis (37, 38), here we identified microbial and host signatures that predict severe disease in both NTM- and NTM+ subjects. Some of these dysbiotic signatures comprised of established pathogens such as *Proteobacteria,* whereas others involved oral facultative anaerobes. By integrating multiple levels of analyses such as biplots, co-occurrence plots and interactomes, we focused on community dynamics rather than one dominant pathogen (39), (20), and discovered an interactive network between oral commensals and NETs in the NTM+ lower airways.

Oral microbes in the lower airways have long been linked with chronic pulmonary inflammation (40–42). In a seminal study, the oral commensal *Neisseria* triggered innate and adaptive immunity and contributed to inflammatory injury in bronchiectasis (43). After identifying microbiota differences in NTM-versus NTM+ BALF, we explored microbial signatures in several phenotypes defined by multiple clinical and radiological measures of severity. Relative abundance of oral commensals was higher in each of the NTM+ severe phenotypes: cavitary, exacerbators and subjects with ‘higher-than-median’ Chest CT scores. Additionally, oral commensals co-occurred with NETs in these NTM+ severe groups. Whether such enrichments and co-occurrences increase the susceptibility to progressive NTM disease remains unknown, but the data presented here supports the paradigm that microbiota beyond a dominant pathogen can contribute to the inflammatory process and disease severity (44).

Targeting neutrophilic inflammation in bronchiectasis has historically led to an increased risk of bacterial infections (45, 46), but neutrophil serine proteases and NETs have emerged as a treatable trait in recent studies. Both the WILLOW and ASPEN trials in bronchiectasis have reported a reduction in exacerbations with Brensocatib, (a dipeptidyl peptidase-1 inhibitor) without compromising neutrophil-mediated bacterial killing. None of these trials however, included NTM+ subjects (7–10). This is justified given that no previous human studies have shown an elevation of NETs in NTM lung infection. Thus, our most novel discovery was that NET was higher NTM+ subjects, and in bronchiectasis subjects with severe disease. NETs also had a synchronous interaction with *Mycobacterium* and oral commensals. The difference in BAL NETs was much more pronounced than that for neutrophil count, suggesting that neutrophil activation, rather than the absolute count may be the more functionally relevant driver *and* marker of NTM+ inflammation. This observation warranted further investigation with a preclinical model. Using our murine model of micro-aspiration (23), we showed a sustained upregulation of NETs, neutrophils, Th-17 and γ8T cells in the NTM+MOC group. Since previous studies have shown that murine NTM-driven immune responses plateau after nine weeks (47), we did not extend our experiments beyond 2 months. Our preclinical findings cannot fully establish a causal directionality between chronic NTM/oral commensals exposure and a NET-inflammatory profile, but it does suggest that experimental reversal of this response could be a potential key step in dissecting mechanistic underpinnings.

Our study has several limitations. Since all samples were collected in a cross-sectional manner, we could not longitudinally evaluate the stability of the microbiome with antibiotics or other treatments. To eliminate the confounding effects of antibiotics, we undertook a subgroup analysis on subjects (74% of NTM-subjects) who had never been treated with any anti-NTM agents and found that the microbiome composition was almost identical to the full cohort. But the effects of NTM treatment *per se* on the pulmonary microbiome remain unknown. Similar to our findings, at least two previous studies did not reveal significant differences in the lung microbiome of bronchiectasis patients who were stable versus those who were on antibiotics for exacerbations (48, 49). Ultimately however, longitudinal studies are required to rule out the effects of different treatment modalities which was outside the scope of our study. Second, given the relatively small proportion of NTM-LD patients with rapid grower sub-species (e.g., *M. abscessus*) we could not conduct sub-analyses but we included the *M. abscessus* patients due to their similar neutrophil and NETosis responses (50–52). Finally, since 16S rRNA gene sequencing lacks strain resolution and functional definition, our future investigations will focus on metagenomic and metatranscriptomic approaches.

In summary, our study represents one of the largest investigations into the lower airway metacommunities and interactomes of bronchiectasis and NTM infection. Using multiple clinical indices of disease, NET assays, clustering and co-occurrence analyses, we described novel dysbiotic signatures in NTM+ bronchiectasis that associate with severe phenotypes. Several of the taxonomic signatures identified include non-dominant pathogens, such as oral commensals. Importantly, these signatures were associated with neutrophilic inflammation and NETs in NTM+ severe disease and in a murine NTM model. Our study, being the first one to sample the lower airways in a large bronchiectasis cohort with a high prevalence of NTM, has the potential to support the exploration of novel NET-targeted therapies for NTM+ patients.

## Authors’ Contributions

LNS and SS conceptualized the study; SS led experiments and wrote the first draft; LNS established the biorepository and finalized the manuscript; FD, KE, IS and BGW did the microbiome analyses; QL and YL helped with the FACS analyses; JKN, MMA and SHC did the co-occurrence analyses; AG, ML and JDC performed the human NET assay; LD, DL and TG helped with the aerosol infection; SG, SP and SK helped with murine NET assay; IA, KK and DA helped with the clinical aspects of the study and biostatistics; SS and EM read the Chest CT scores; SK, SP, CC, YL, MC, AH, RS, AL, SG, KW, BK, CB, TK, IK, RS, CR, DH, GP, YF, AC, CM, EC, YK, MK, JT, JN, NN, AB, DK, MW and DAH helped with all clinical aspects of the study and other analyses.

## Research Support Funding

R37 CA244775 (LNS, NCI/NIH); U2C CA271890 (LNS, NCI/NIH); U01 AG088351 (LNS, NIA/NIH); R33 GM147800 (LNS, NIH/NIGMS); American Thoracic Society Bronchiectasis Initiative Award (SS); Doris Duke-FRCS Award (SS); CTSI KL2 Scholar Award (SS); Stony Wold-Herbert Fund Grant-in-Aid/Fellowship (SS); Veterans Affairs IK2BX005309-01A2 (BGW); CHEST Foundation Research Grant in Chronic Obstructive Pulmonary Disease (BGW). The Genome Technology Center is partially supported by the Cancer Center Support Grant P30CA016087 at the Laura and Isaac Perlmutter Cancer Center (AH, AT)

## Scientific Knowledge on the Subject

Lower airway dysbiosis and *Pseudomonas* dominance have historically been associated with aberrant neutrophil function and severe phenotypes in bronchiectasis. Whether such airway dysbiotic signatures are associated with neutrophil extracellular trap formation and disease progression/severity in NTM+ bronchiectasis is not known.

## What this Study Adds to the Field

We identified that lower airway dysbiosis is associated with severe endophenotypes in bronchiectasis, including NTM positivity and elevated neutrophil extracellular trap levels. Mycobacterial infection and oral commensals both contributed to a Th17-neutrophilic immune response and drove neutrophil extracellular trap formation in a murine micro-aspiration model.

## Additional Information

S.H.C serves on the advisory boards for CSL Behring, Sanofi, GSK, Zaccha Pte Ltd. Pneumagen Ltd. and Boehringer Ingelheim; has received lecture fees from AstraZeneca, CSL Behring, Boehringer-Ingelheim and Chiesi Farmaceutici and has served on Data Safety and Monitoring Boards for Inovio Pharmaceuticals Ltd. and Imam Abdulrahman Bin Faisal University. Other authors have no competing financial or non-financial interests to disclose.

## Methods

### Study Design and Subject Recruitment

Our study was a prospective observational study of 200 patients with a diagnosis of non-cystic fibrosis bronchiectasis enrolled over 7 years (2014–2021). All subjects were enrolled from a bronchiectasis cohort at New York University (NYU) Langone Medical Center and signed an informed consent form to participate in the study. The research protocol was approved by the NYU Institutional Review Boards (IRB# S14-01400). The study inclusion criteria included Chest CT imaging abnormalities and symptoms consistent with bronchiectasis. Exclusion criteria included participants who had been on antibiotics for an infectious exacerbation with organisms other than NTM within the last month, thus excluding subjects who had been on antibiotic regimens commonly used in bronchiectasis subjects such as macrolides, fluoroquinolones and cephalosporins. Patients with acute pneumonias, concurrent malignancy, and those on chronic therapies such as immunoglobulins, suppressive macrolide therapy, steroids and biologics were also excluded. At the time of recruitment, questionnaire data (St. George’s Respiratory Questionnaire and Reflux Symptom Index) was obtained.

### Clinical Data Indices

Demographic and clinical information was collected via review of electronic medical records. Patient characteristics were collected at enrolment and included age, sex, race, body mass index (BMI), smoking status, history of other co-morbidities, lung physiology data including spirometric volumes/ lung volumes/ diffusion coefficient, FACED scores, number of bronchiectasis exacerbations in the last year, radiological features, and culture results for secondary pathogens such as *Pseudomonas aeruginosa, Aspergillus spp.,* and *Haemophilus influenzae.* The severity of bronchiectasis was evaluated using the FACED Score (1). FACED stands for **F**EV1% predicted, **A**ge, **C**hronic colonization (specifically with *Pseudomonas aeruginosa*), radiological **E**xtent, and **D**yspnea score (as per the modified Medical Research Council score). The score assigns points based on these five variables, resulting in a total score ranging from 0 to 7: Mild: 0-2 points, Moderate: 3-4 points and Severe: 5-7 points. Exacerbations were recorded based on the consensus definition for bronchiectasis exacerbations (2) (3). Patients were stratified into groups based on the frequency of exacerbations in the one year prior to enrolment. For the purpose of this study, “non-exacerbators” had 0-1 exacerbations in the preceding year and “exacerbators” had >/= 2 exacerbations in the preceding year (2) (3). The radiological groups of cavitary versus non-cavitary disease was based on the presence or absence of lesions that were visualized as a gas-filled space within a nodule or area of consolidation. Fibrocaseous features that are marked by fibrosis and honeycomb cysts were not reported as cavities. Finally, a detailed Chest CT score was used to delineate the extent and severity of bronchiectasis and NTM-LD, as below.

### Scoring of Chest CT Scans

All subjects had a Chest CT scan done prior to bronchoscopy. Chest CT scans were scored on the basis of the following three criteria:

a) <u>Reiff Score for Bronchiectasis:</u> Modified from Bhalla’s (4) Chest CT scoring system for Cystic Fibrosis, the Reiff score (5) is now commonly used for delineating the radiological disease burden in non-cystic fibrosis bronchiectasis. The Reiff Score has been refined over the years to score on the basis of tubular, varicose and cystic changes in the six lung lobes (6) (7) (8). It allows for one point for the presence of tubular bronchiectasis, two points for the presence of varicose bronchiectasis and three points for the presence of cystic bronchiectasis in any particular lobe. The maximum score possible under this criterion is 18.
b) <u>Size and number of cavities:</u> Cavities in all six lobes were scored on the basis of size and number, as previously described for cystic fibrosis and NTM-LD (9) (10). One point was scored for every 1cm increment in the size of cavities, up to a maximum of 9cm (nine points). One point was also given for n=1 of cavities, up to a maximum of n=9 (nine points). The maximum score possible under this criterion is 18.
c) <u>Presence of tree-in-bud and nodules:</u> One point was scored for the presence of nodules or tree-in-bud changes in each of the six lobes (9) (10). The maximum score possible under this criterion is 6.

The maximum possible Chest CT score was 42. In our cohort, the median Chest CT score was 13 [IQR 12-16], median cavity size was 2.4 cm [IQR 2-4] and median number of cavities was 1 [IQR 1-4.25].

### Diagnosis of NTM-LD

Diagnosis of NTM-LD was made in accordance with the American Thoracic Society diagnostic criteria (11). Out of the 92 subjects who were NTM+, 73/92 (79%) had been diagnosed based on a positive bronchoalveolar lavage fluid (BALF). One patient had an NTM+ transbronchial biopsy. The remaining 18/92 (20%) subjects were diagnosed on the basis of two NTM positive sputum cultures that had all been collected within 60 days of the date of the bronchoscopic sampling.

### Bronchoscopy and sample collection

As per our protocol, we asked every patient enrolled in the cohort about their interest in participating in a bronchoscopy arm. A few patients agreed to a research bronchoscopy (n=6). Other 194 subjects had a bronchoscopy done as per clinical indication (in general this was because of difficulties with obtaining three induced sputum samples or the persistence of clinical symptoms suspicious of NTM infection) and patients agreed to have bronchoscopic samples obtained for this research. BAL samples were collected from the radiologically most diseased lung segment in every subject, as confirmed by the Chest CT score (above). For all the BAL sampling, we used room temperature normal saline. Three aliquots of 50 mL each were injected and withdrawn 3 times, with careful measurement of volume in and volume out, as per standard practice. Mild suction was applied to prevent airway collapse, with intermittent suctioning. In all subjects who underwent bronchoscopy, we also collected supraglottic samples (SUP) and background/ equipment samples (BKG). SUP samples were collected from the throat/supraglottic region using a Yankauer and BKG samples were collected as a saline flush through the bronchoscope prior to the bronchoscopy.

### Treatment History for NTM-LD

In a subgroup analysis, we excluded 15/92 (16%) subjects who were on NTM treatment at the time of the bronchoscopy as well as 9/92 (10%) subjects who had treatment in the past. Therefore, for this subgroup analysis, we focused on 68/92 (74%) of the NTM+ subjects who had never been treated with any of the first/second/third line agents against NTM. We refer to this sub-group as never-treated NTM+ subjects.

### Lower airway bacterial load and taxonomy using 16S rRNA gene sequencing

For DNA extraction, we used the Qiagen DNA mini kit. The bacterial load was measured using a QX200 Droplet Digital PCR (ddPCR) System (BioRad, Hercules, CA) (12). Cycling conditions included 1 cycle at 95°C for 5 minutes, 40 cycles at 95°C for 15 seconds and 60°C for 1 minute, 1 cycle at 4°C for 5 minutes, and 1 cycle at 90°C for 5 minutes all at a ramp rate of 2°C/second. High-throughput sequencing of bacterial 16S rRNA-encoding gene amplicons (V4 region) was performed on all upper airway and BAL samples. In addition, background control samples (DNA free water through suctioning the channel of the bronchoscope obtained prior to bronchoscopy) were also sequenced and analyzed in parallel. Barcoded 16S rRNA gene amplicons encoding the V4 region were generated (13). Amplicons were quantified using Agilent 2200 TapeStation system and then pooled for sequencing on the Illumina MiSeq (150bp read length, paired-end protocol) in one single run. The 16S rRNA gene sequences obtained were analyzed using the Quantitative Insights into Microbial Ecology 2 (QIIME2) 2-2022.8 package. Sequence reads were demultiplexed with quality control (14). DADA2 workflow was then applied and amplicon sequence variants (ASVs) were obtained. ASVs were then assigned taxonomy using Greengenes 99% ASVs (15). Resulting 16S rRNA gene data was used for further analysis in *R* (version 4.1.0) using functions provided with QIIME2R (16) (version 0.99.6), Phyloseq (17) (version 1.38), Vegan (version 2.6-2) and Tidyverse (version 1.3.2). Microbiome diversity indices were evaluated by comparing alpha diversity measures (Richness, Shannon index and Simpson index); and beta diversity principal coordinate analyses (PCoA) based on the Bray-Curtis dissimilarity index. Differential taxonomic enrichment data between different groups was performed using EdgeR package with an FDR cut-off of 0.2 (version 3.36.0) (18–20). We applied Dirichlet multinomial mixtures (DMM) modeling on all lower airway samples to identify distinct profiles of lower airway microbiota (21).

### Contaminomics

Since low biomass samples can be significantly impacted by the presence of contaminants, a prevalence-based method from the *decontam* package in *R* (version 1.14.0) was used (22). In this process, all reads from lower airway samples were compared against background controls. Taxa identified as potential contaminants were not removed from subsequent analysis, given the compositional nature of the data, rather their label as a potential contaminant was retained and referenced throughout all downstream analyses. All contaminants are listed in Table E1 and Figures E2A-C.

### Microbial topographical analysis based on pathogenic potential

In order to determine the abundance of pathogenic taxa in each sample, all taxa underwent a process of sorting by pathogenic potential. The purpose of this analysis was to identify patients whose lower airway samples demonstrated a high relative abundance of pathogenic taxa by 16S sequencing (as opposed to culture-based pathogen identification alone) and assess whether these pathogen-rich samples were associated with differences in microbiome diversity and clinical phenotype. Taxa were excluded from this analysis if they did not comprise at least 0.1% relative abundance in any single sample (excluding 2,492 of 7,518 taxa). Pathogenic potential classification was assigned based on each taxon’s genus because species-level taxonomic identification was often not available via QIIME2 and targeted 16S rRNA sequencing data lack the necessary precision to accurately label species. However, 1,821 taxa did not have a genus-level taxonomy available and were excluded from the analysis. In order to avoid excluding important taxa, those with at least 1% relative abundance in at least one sample underwent NCBI BLAST database search, resulting in genus identification and inclusion of 302 of these 397 taxa. A total of 3,507 taxa were then individually searched on Pubmed to assess for reported respiratory infections, then categorized into one of three pathogen categories: Pathogen (P), Possible Pathogen (PP), or Non-Pathogen (NP). A genus was defined as pathogenic if it contained pathogens that commonly cause pneumonia in immunocompetent persons. A genus was defined as a possible pathogen if a Pubmed search returned >10 reports of infection in humans, regardless of it being classified as an opportunistic infection. If a Pubmed search returned a few isolated case reports of human infection or no reports at all, the genus was defined as a non-pathogen. Once all taxa were categorized as pathogenic or not, the percentage composition of pathogenic, possible pathogenic, and non-pathogenic taxa in each patient sample was determined. Samples composed of >50% relative abundance belonging to pathogenic taxa (sum of the genera annotated as pathogenic) were defined as pathogen rich. Samples composed of >50% relative abundance belonging to possible pathogens (sum of the genera annotated as possible pathogens) were defined as possible pathogen rich. Table E3 shows the classification of all taxa into pathogenic, possible pathogenic or non-pathogenic species.

### Human BAL neutrophil extracellular trap (NET) immunoassay

There is no agreed upon high-throughput method of quantifying NETs in human biological fluids. The human BAL NET immunoassay was done in Dr James Chalmers laboratory at the University of Dundee, UK as a collaborative effort. Dr. Chalmer’s laboratory has previously published this immunoassay in several seminal research manuscripts (23, 24). The assay has been developed and validated in house and is based on detection of DNA-elastase and histone-elastase complexes. For the DNA-elastase complex assay, anti-DNA (HYB331-01; Abcam, Cambridge, United Kingdom) capture antibody was incubated on plates overnight at 48C, followed by washing with PBS plus 0.05% Tween 20 (wash buffer). Plates were blocked with 1% BSA in PBS and washed with wash buffer. Samples were diluted in 1% BSA in PBS. The BAL samples were first diluted 1 in 5 and if high, we then to a 1 in 50 or in some cases a 1 in 100 dilution to get values in the dynamic range/within the standard curve. The standard curve was generated by titrating concentrations of heathy human blood-derived neutrophils treated with phorbol 12-myristate 13-acetate (PMA). Plates were washed 3 times with wash buffer after incubation of standards and samples. DNA-elastase complexes were detected with sheep anti–neutrophil elastase–horseradish peroxidase (PA1-74133; Thermo Scientific, Waltham, Mass) and developed with 3,395,59-tetramethylbenzidine. For the histone-elastase assay, plates were coated for 1 hour with anti–histone H1 (ab71594, Abcam), washed and blocked as above, and incubated for 1 hour with rabbit anti–neutrophil elastase (ab21595, Abcam). Anti-rabbit–horseradish peroxidase (ab6721, Abcam) was used for detection, and the plate was developed as above. The assays were validated against other known NET components (citrullinated histone H3 and DNA) for the effects of sample preparation methods and for passive interactions between DNA and elastase.

### Urea Dilution Factor to standardize BAL NET analysis

The urea dilution method was used to standardize the concentration of NETs in the BALF. Plasma urea/ BAL urea ratio was calculated for each sample. The ratio was then used as a dilution factor by which the NET concentrations were adjusted. BAL urea level was calculated using the Invitrogen blood urea nitrogen colorimetric detection kit (EIABUN), as per manufacturer’s instructions. The corrected BAL NET levels have also been reported in Table 1.

### Human BAL cell count

Acellular bronchoscopy samples were used for the measurement of cell count and differentials as previously described (13, 25).

### Co-occurrence analyses

In order to evaluate microbe-microbe associations and/or microbe-cell type associations, co-occurrence analyses were performed. For these analyses, all microbial counts were aggregated at genus level before more downstream analysis. Only microbes with greater than 1% abundance in at least 5% (n=10) of the patients were retained. Subsequently, the filtered microbiome datasets and inflammatory cell counts were normalized to percentages for further analysis. Co-occurrence analysis networks were derived using an ensemble correlation-based (Spearman and Pearson) network inference algorithm coupled to ReBoot (Bootstrap with renormalization) mitigated against compositionality adapted from previous studies (26) (27). Clinical subgroups analyzed were NTM negative, NTM positive, exacerbators and subjects with cavitary disease. To derive microbiome and inflammatory cell associations networks, microbiome and cell counts data were normalized to percentages after concatenation. Edge weights were corrected for multiple comparisons using the Benjamini and Hochberg method FDR (False Discovery Rate) procedure and only edges with a corrected p-value < 0.001 were considered and reported (26). More details of the network inference algorithm have been described in previous publications (28). Nodes (microbes and/or cells) within the network were ranked based on their interaction ‘activity’ defined as <u>a)</u> busy: degree of a node - microbes with an increased number of direct interactions with other microbes; <u>b)</u> critical: stress centrality - key microbes to maintain network integrity; and <u>c)</u> influential: betweenness centrality - microbes influencing other microbes in a network, including indirectly. Red lines demonstrate a negative or competitive interaction (negative correlation – an increase in one microbe’s abundance correlates with a decrease in another’s) whereas green lines demonstrate a positive interaction (positive correlation – an increase in one microbe’s abundance correlates with an increase in another’s). The highest calculated network metrics are highlighted by size, width and node coloration, respectively, in the presented network plots. All the networks were plotted and visualized using R and Cytoscape.

### Microbiome-NET association analysis

To evaluate the relationship between compositional microbiome and non-compositional NET data, a linear model within the MaAsLin2 (Microbiome Multivariate Association with Linear Models) framework (29) was implemented within each clinical subgroup (see above). The microbiome dataset underwent normalization using CLR (centred-log-ratio transformation), while NET values were standardized with z-scaling prior to MaAsLin analysis using Linear models. To ensure robust results, statistical comparisons within the MaAsLin framework were adjusted for multiple testing using the Benjamin-Hochberg method. Associations with q-values <0.1 were considered statistically significant. The resultant association are visualized as network plots using networkx-package (python), with edge weights representing the model coefficient (strength of association).

### Animals-mice

Wild type BALB/c mice (6-8 weeks of age) were purchased from Jackson Research Laboratories, Bar Harbor, ME Cat#000651. Recent studies have shown BALB/c mice to be a good model of chronic NTM infection when compared to C57BL/6 or the C3HeB/FeJ strains(30),(31). This was confirmed by us in optimization experiments using all three strains. Mice were allowed 2 weeks of acclimatization to the facilities prior to the start of experiments. M ice with different experimental conditions were co-housed to limit potential cage effect on the microbiome and host immune tone. Mice received sterilized food and water *ad libitum*.

### Human oral commensals

#### Composition

We used the human oral commensals *Veillonella parvula* (ATCC 17742), *Prevotella melaninogenica* (ATCC 25845) and *Streptococcus mitis* (ATCC 49456) for these experiments. This combination of the oral commensals was chosen based on the following seminal studies:

a) A study from us [Segal et al (13)] that showed that enrichment with these oral taxa (referred to as Pneumotype_SPT_) was associated with a distinct metabolic profile and a pro-inflammatory Th17 phenotype.
b) A study of integrative microbiomics in bronchiectasis exacerbations that identified these oral taxa in high exacerbation frequency clusters (28).
c) Data from the Bronchiectasis and Low-Dose Erythromycin Study (BLESS) that showed that bronchiectasis patients with more exacerbations had a *Veillonella* dominated microbiome (32).

#### Dose

The dose of the oral commensals used in this experiment was based on previous work and publications from our lab that investigated single versus multiple challenges and dose-response experiments using low, medium and high concentrations of the bacteria (33). Based on these previous studies, we aimed for 10^9^ CFUs/ml in the inoculum for each of the bacterium. Considering all experiments, the median concentration of *Veillonella Parvula* used in our experiments was 7.87×10^9^ /ml, the median concentration of *Prevotella melaninogenica* used was 2.36×10^9^ /ml and the median concentration of *Streptococcus mitis* was 1.94×10^9^ /ml. Each of the bacteria were mixed in a 1:1:1 ratio and the aliquots used for intra-tracheal challenge have been referred to as a Mixture of Oral Commensals (referred to as MOC and consisting of *Streptococcus*, *Veillonella* and *Prevotella*). MOC was used in 50 µL aliquots. As in previous studies (34), none of the oral commensals, since they were strict anaerobes, could be retrieved post-inoculation, from lung tissue homogenates.

#### Anaerobic chamber

Bacteria were grown in anaerobic conditions (Bactron 300, Shel Labs, Cornelius, OR), then stored in 20% glycerol tryptic soy broth at -80°C. To prepare the oral commensal challenges, bacterial strains were thawed and streaked on anaerobic PRAS-Brucella Blood agar plates (Anaerobe Systems, Morgan Hill, CA). Plates were incubated at 37°C in an oxygen-free environment (tri-mix: 5% carbon dioxide, 5% hydrogen and 90% nitrogen) in an anaerobic chamber for 24-48 hours. colonies were collected from the plate and re-suspended in 1 ml of sterile PBS. OD620 was measured to calculate the approximate concentration prior to use.

### Murine experimental design

WT BALB/c mice were divided into four experimental groups: <u>Group 1:</u> No intervention (Untreated), <u>Group 2:</u> NTM aerosol challenge only (NTM), <u>Group 3:</u> Weekly intratracheal MOC challenge (MOC), and <u>Group D:</u> NTM aerosol challenge followed by weekly MOC (Fig. 6A). MOC was delivered on a once-weekly basis. All mice were sacrificed either after one month or two months after the day of the NTM infection. Each group had 5 mice. We have performed these experiments at least six times at different time points.

### Murine aerosol NTM infection

BALB/c mice were infected with the MAC 101 NTM strain (kind gift from Dr. Eric Neurmberger, JHU) via aerosol delivery. The infecting inoculum was prepared from freshly growing cultures after three washes, at a concentration of approximately 10^8^ CFU/ml. The nebulization was undertaken using an Aeroneb lab nebulizer system from Kent Scientific (https://www.kentscientific.com/products/aeroneb-lab-nebulizer-unit/). This system allows a particle size of 2.5μm to 4μm, has a flow rate of >0.1 ml/min and a residual volume of <0.1ml. The nebulizer system was connected to an oversized box that would hold up to 30 mice for each round of nebulization. Final volume nebulized was approximately 5mls over 30 minutes. Mouse lung NTM CFU’s were consistently in the range of 10^7^ CFU/ml for several weeks after infection, thus confirming a bacterial deposition rate of 10%. This is comparable to the Glas-Col nebulization system used in BSL-3 laboratories. Mice were awake during the nebulization.

### Intratracheal MOC inoculation

For the intratracheal MOC inoculation, mice were anesthetized utilizing isoflurane via VetFlo Anesthesia machine (Kent Scientific, Torrington, CT) sedated to 10-15 breaths per minute and monitored for any distress. Mice were then placed on an intubation platform and with blunt forceps, their tongue was gently pulled ventrally until the pharynx was exposed. A human otoscope with a 2 mm ear cone (Welch Allyn 3.5V Hill-Rom, Inc., Skaneateles Falls, NY Model #20200) was introduced into the oral airway to expose to visualize the murine vocal cords and a gel-loading pipette tip loaded with a 50μL aliquot was introduced through the vocal cords of the mouse and deployed into the lower airway. The mouse was removed from the platform to recover from anesthesia on a heat pad. Mice were monitored every 2-4 hours following intra-tracheal challenge.

### FACS analysis

Fluorescence-activated cell sorting (FACS)/ flow cytometry was undertaken with single cell suspensions derived from lung homogenates as previously described by us (34). Briefly, single-cell suspensions were prepared from lung ∼1-mm lung segments, followed with incubation at 37°C for 30 min in RPMI 1640 in the presence of Liberase TM (50 μg/ml, Roche) and DNase I (1μg/ml, Thermo Fisher Scientific). Digested lung tissues were then mashed through 70μm filters into complete RPMI-1640 with 10% fetal calf serum. A 30:70% (v/v) isotonic discontinuous Percoll (GE Healthcare) density gradient was used for mononuclear cell enrichment. Cell viability (>95%) was determined by trypan blue exclusion. Fluorescence-conjugated Abs for surface staining, including anti-CD45.2 (104), anti-CD4 (RM4-5), anti-CD8α (53-6.7), anti-γδTCR (GL3), anti-CD11b (M1/70), anti-CX3CR1 (SA011F11), anti–Ly-6C (AL-21), anti–Ly-6G (1A8), anti-Siglec-F (S17007L), anti-CD11c (HL3), anti-MHC II (M5/114.15.2), anti-PD-1 (RMPI-14), anti–IL-17A (TC11-18H10.1), and anti-RORγt (Q31-378) were purchased from BioLegend (San Diego, CA), eBioscience/Thermo Fisher Scientific (Waltham, MA), or BD Biosciences (San Jose, CA). Foxp3 (PJK-16s; eBioscience) was quantified by intracellular staining performed according to the manufacturer’s protocol. To detect the expression of IFN-γ, Th17 cells were stimulated for 4–5 h with PMA (Sigma-Aldrich, St. Louis, MO) and ionomycin (Sigma-Aldrich) in the presence of Brefeldin A (Sigma-Aldrich) and then stained with anti–IFN-γ (XMG1.2) and anti-IL-17A mAb. Flow cytometry was performed with BD FACS Symphony A5 SE (BD Biosciences) and analysis was performed with FlowJo software (Tree Star).

### Murine BALF NET ELISA

Semiquantitative ELISA-based detection of DNA-H3Cit complexes was performed as previously described (35). 50 µL of a 2.5-µg/mL solution of murine monoclonal antibody against anti-double stranded DNA (cat. no. ab27156, Abcam) in PBS was added to each well of a high-binding 96-well plate (cat. no. 3369, Corning) and incubated overnight at 4°C. The plate was washed 3 times with 400 µL PBS per well, blocked with 200 µL of 1% BSA in PBS per well for 2h at room temperature, and washed again. Then, 100 µL of BAL supernatant (collected by centrifuging BAL for 2 min at 2,000 *g*) was added to each well and incubated for 2 h at room temperature. Wells were washed twice, and 100 µL/well of a 1-µg/mL solution of anti-H3Cit detection antibody (cat. no. ab5103, Abcam) in PBS was added before incubation for 1 h at room temperature. After being washed twice, 100 µL/well of anti-rabbit horseradish peroxidase-conjugated antibody (1:1,000 dilution; cat. no. 7074, Cell Signaling Technology) was added and incubated for 1 h at room temperature. Three more washes were performed, 100 µL of TMB solution (cat. no. 34028, ThermoFisher) was added to each well, and the reaction was incubated for 10 –20 min at room temperature on a rotary shaker, avoiding direct light. We added 50 µL of 2N HCl to each well to stop the reaction. The plate was then read immediately at 450 nm and 560 nm (a reduction measurement to correct for background) using a plate reader (Cytation 3, BioTek). Technical triplicates were used for each sample and the experiment was repeated three times.

### Statistical analysis

Since the distributions of microbiome data are non-normal, no distribution-specific tests are available. Therefore, we used non-parametric tests of association. We used either the Mann-Whitney test or the Kruskal-Wallis ANOVA (in case of > 2 categories). Wilcoxon signed-rank test were used for paired analysis. To cluster microbiome communities into exclusive ‘metacommunities’ we used the Dirichlet Multinomial Mixture Model (21). In this method, for each sample, we impute the component most likely to have generated it, thus separating samples into groups it has the highest probability of belonging to. This allows for variable cluster sizes and a more rigorous means of choosing optimal cluster number. To evaluate differences between groups of 16S data, we used linear discriminant analysis (LDA) Effect Size (LEfSe) (36). For tests of association with continuous variables, we used non-parametric Spearman correlation tests and FDR was used to control for multiple testing.

### Data availability

Sequencing data are available in NCBI’s Sequence Read Archive under project number PRJNA1049425. Codes utilized for the analyses in the manuscript are available at https://github.com/segalmicrobiomelab/Bronchiectasis_lower.airway_microbiome_paper.

## Supplementary Tables

**Table E1.** We evaluated for potential bacterial DNA contaminants in our dataset using Decontam. Given the compositional nature of the data and challenges in confirming the “contaminant” nature of the taxa identified, we did not remove these taxa from subsequent analysis. Instead, we labelled them as potential contaminants for reference.

**Table E2.** Blast analyses of the most abundant *Mycobacterium* ASVs identified *Mycobacterium avium* as the top potential strain. For the other four NTM-subjects with *Mycobacterium* reads at low abundance, blast analyses of the ASVs showed that the top annotations were consistent with non-pathogenic *Mycobacteria* (such as *Mycobacterium frederiksbergense, Mycobacterium hemophilum* and *Mycobacterium lacus*) rather than pathogenic *Mycobacteria*.

**Table E3** shows the classification of all taxa into pathogenic, possible pathogenic or non-pathogenic species.

## Supplementary Figures

**Figure E1.**
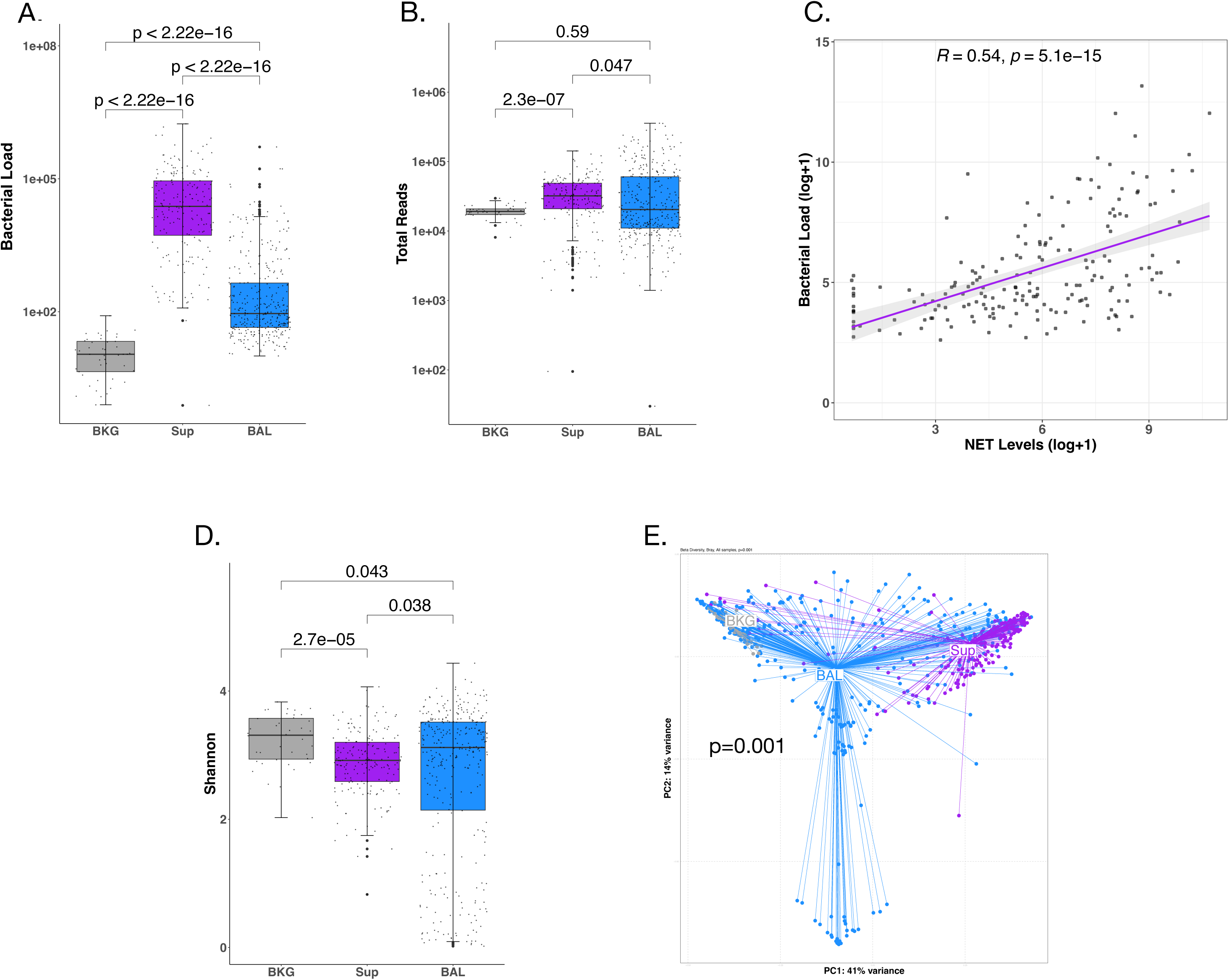
Evaluation of the lower airway microbiota. Fig. E1A. Bacterial load (ddPCR) was lowest in BKG samples, followed by BAL samples and highest in Sup samples (p<0.0001 for all comparisons, Mann Whitney). **Fig. E1B.** Total reads were also lowest in BKG samples, followed by BAL samples and highest in Sup samples (BKG versus Sup p<0.001, Mann Whitney). **Fig. E1C.** Spearman’s rank correlation coefficient between BAL bacterial load and NET levels was highly significant, regardless of NTM status (Spearman’s p= 5.08e-15). **Fig. E1D.** Alpha diversity as measured by Shannon index was lower in the Sup samples than in BAL samples (p=0.038, Mann Whitney). **Fig. E1E.** Bray Curtis Dissimilarity index showed clear compositional differences between the three sample types (p=0.001, PERMANOVA).

**Figure E2.**
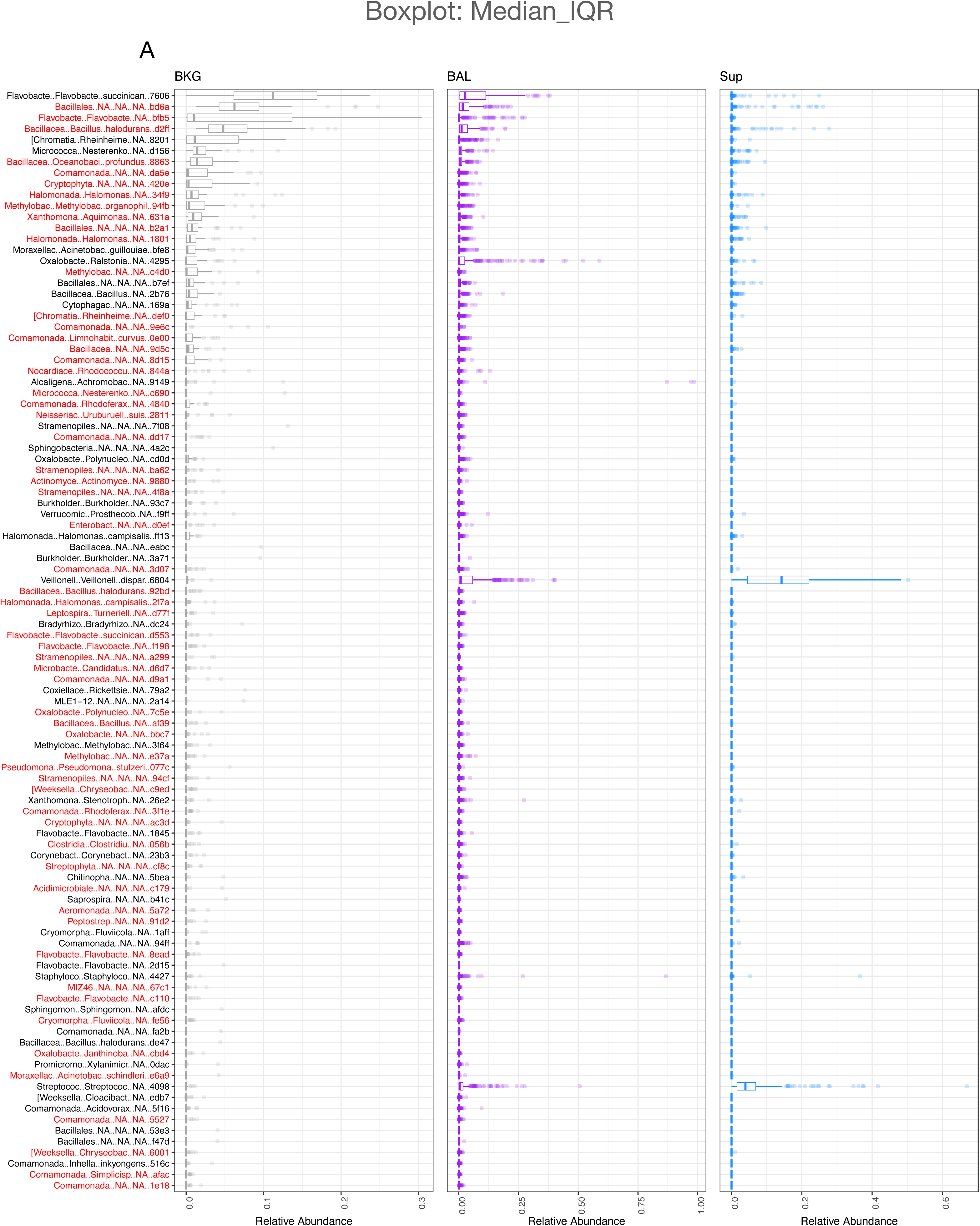
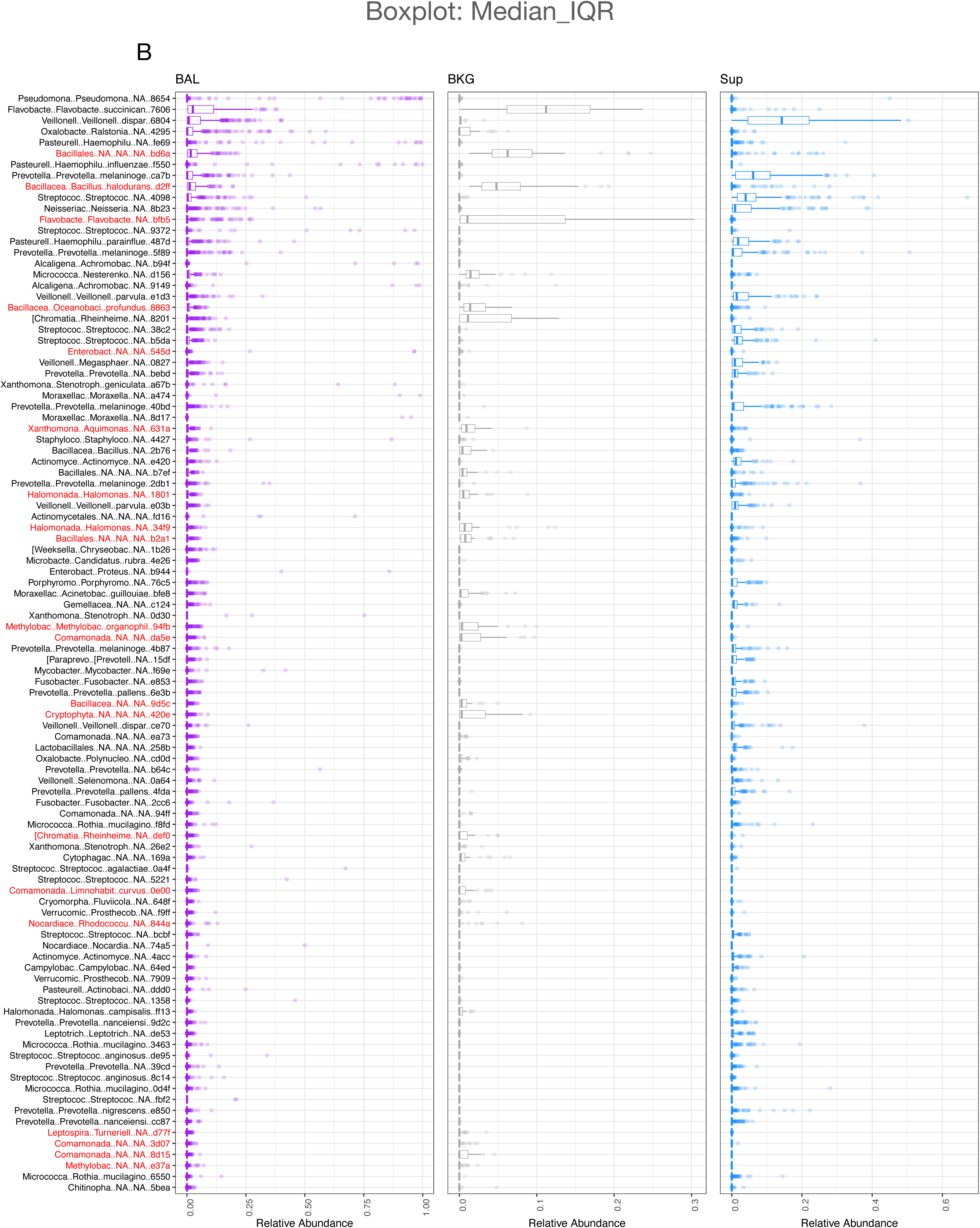
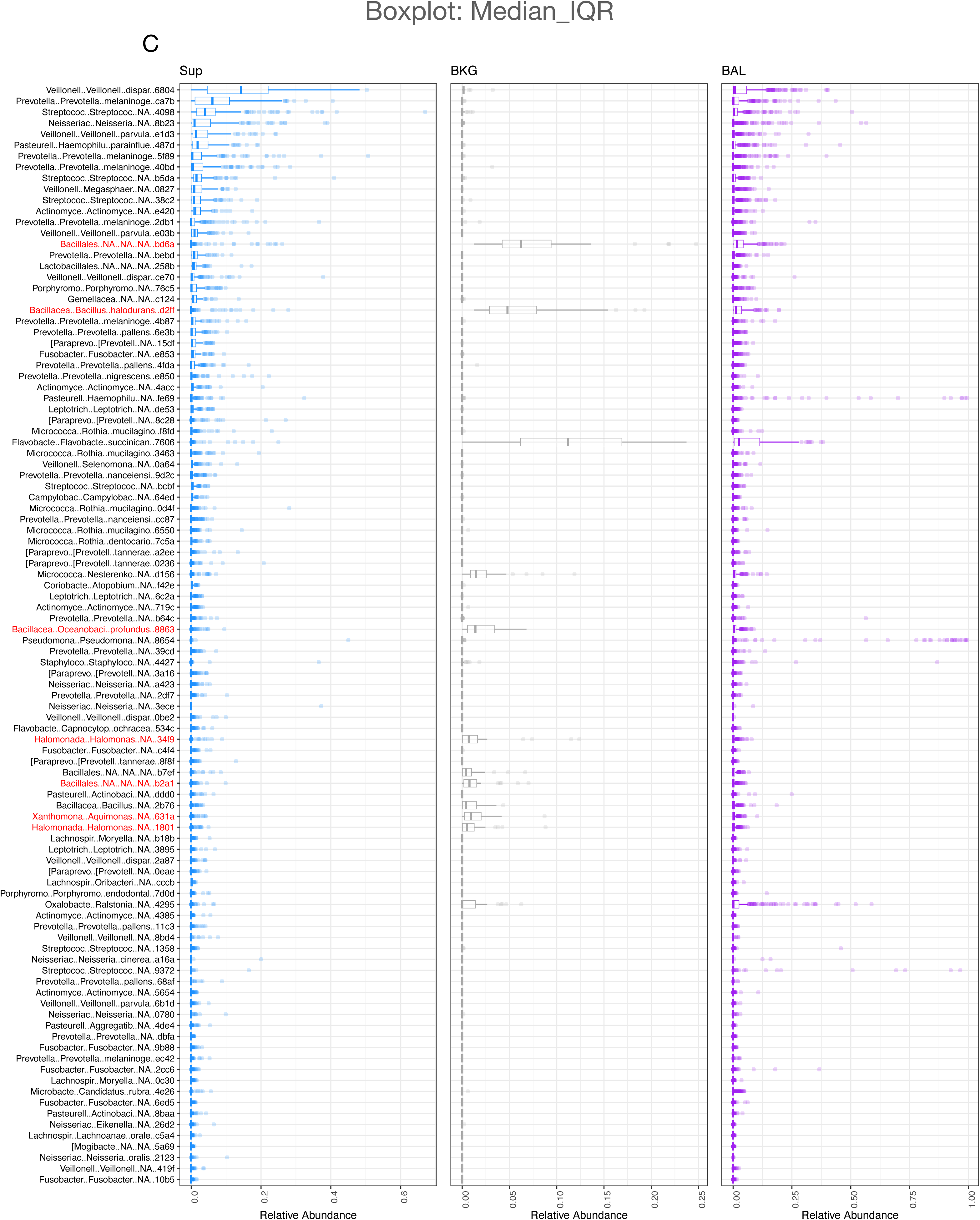
Evaluation for potential bacterial DNA contaminants with *Decontam.* Fig. E2A. shows the median and IQR for all the potential bacterial DNA contaminants (red font) in the BKG samples, **Fig. E2B.** in the BAL samples and **Fig. E2C.** in the Sup samples. We compared the relative abundance of taxa in BKG samples to Sup and BAL samples to identify potential contaminants by the prevalence method. A total of 195 out of 7518 taxa were identified as potential contaminants and the top potential contaminants included *Bacillales* and *Flavobacterium*.

**Figure E3.**
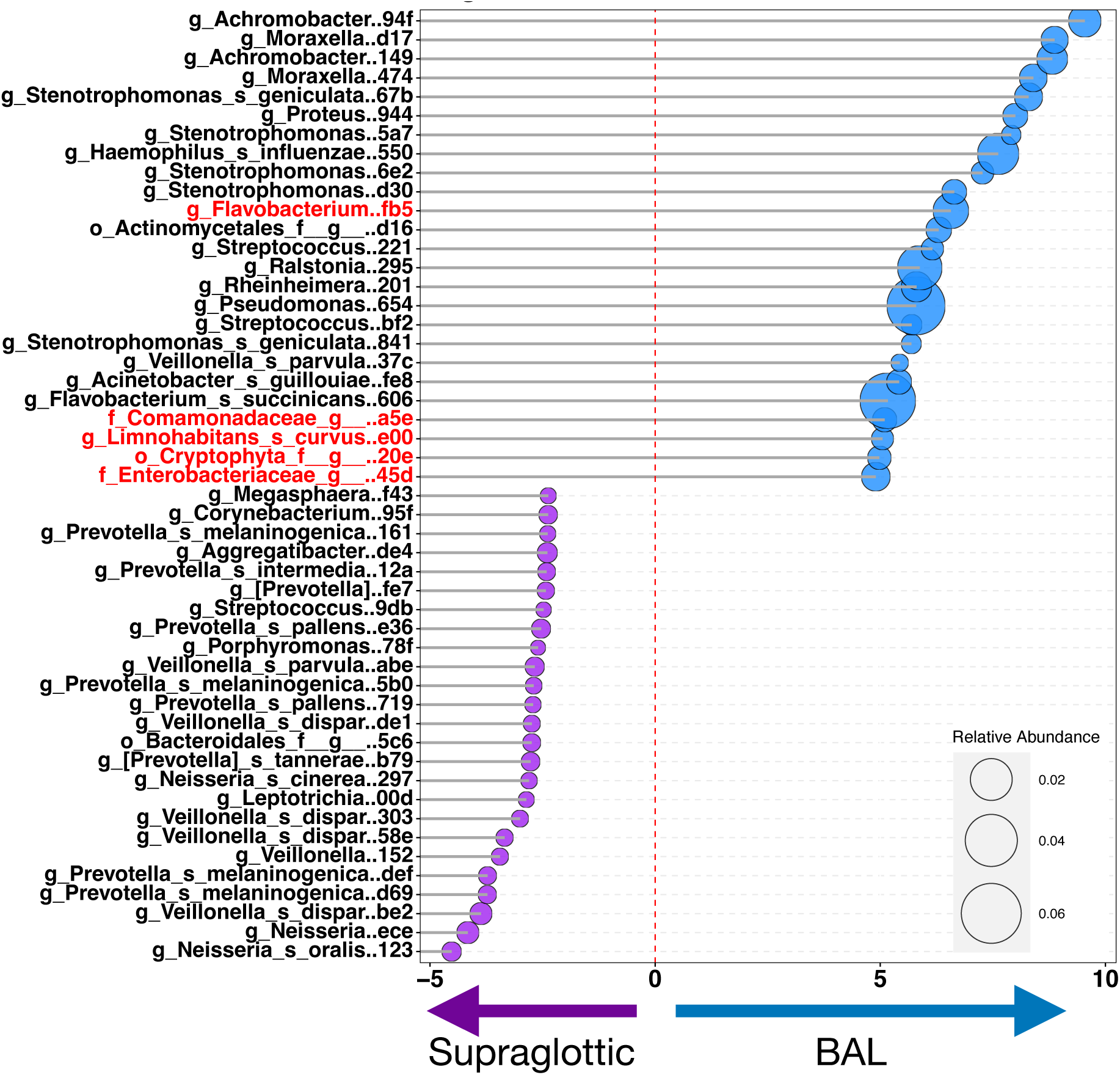
Differential taxonomic enrichment between upper and lower airways. EdgeR analysis revealed that BAL samples had a higher relative abundance of *Hemophilus, Ralstonia, Flavobacterium* and *Pseudomonas* when compared to Sup samples.

**Figure E4.**
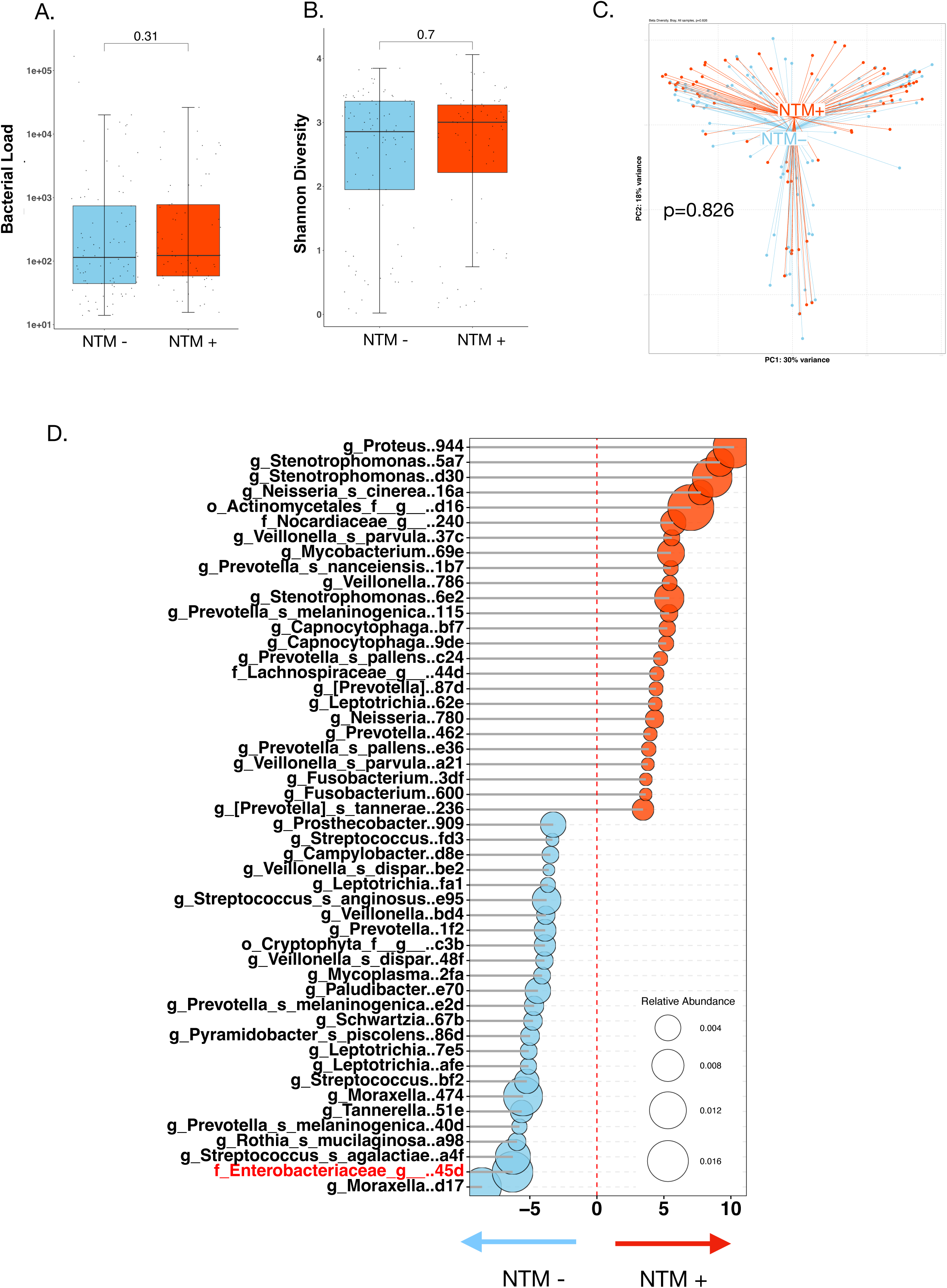
Comparison of the lower airway microbiota in never-treated NTM-/ NTM+ subjects. Fig. E4A-C. Similar to the full cohort, there were no significant differences in the bacterial burden, alpha diversity or beta diversity between NTM- and NTM+ never-treated subjects. **Fig. E4D.** Taxonomically, the never-treated NTM-BAL was noted to have a higher relative abundance of *Moraxella, Enterobacteriaceae, Streptococcus* and *Actinobacillus,* and the never-treated NTM+ BAL had a higher relative abundance of *Proteus, Stenotrophomonas, Actinomycetes* and *Nocardia*.

**Figure E5.**
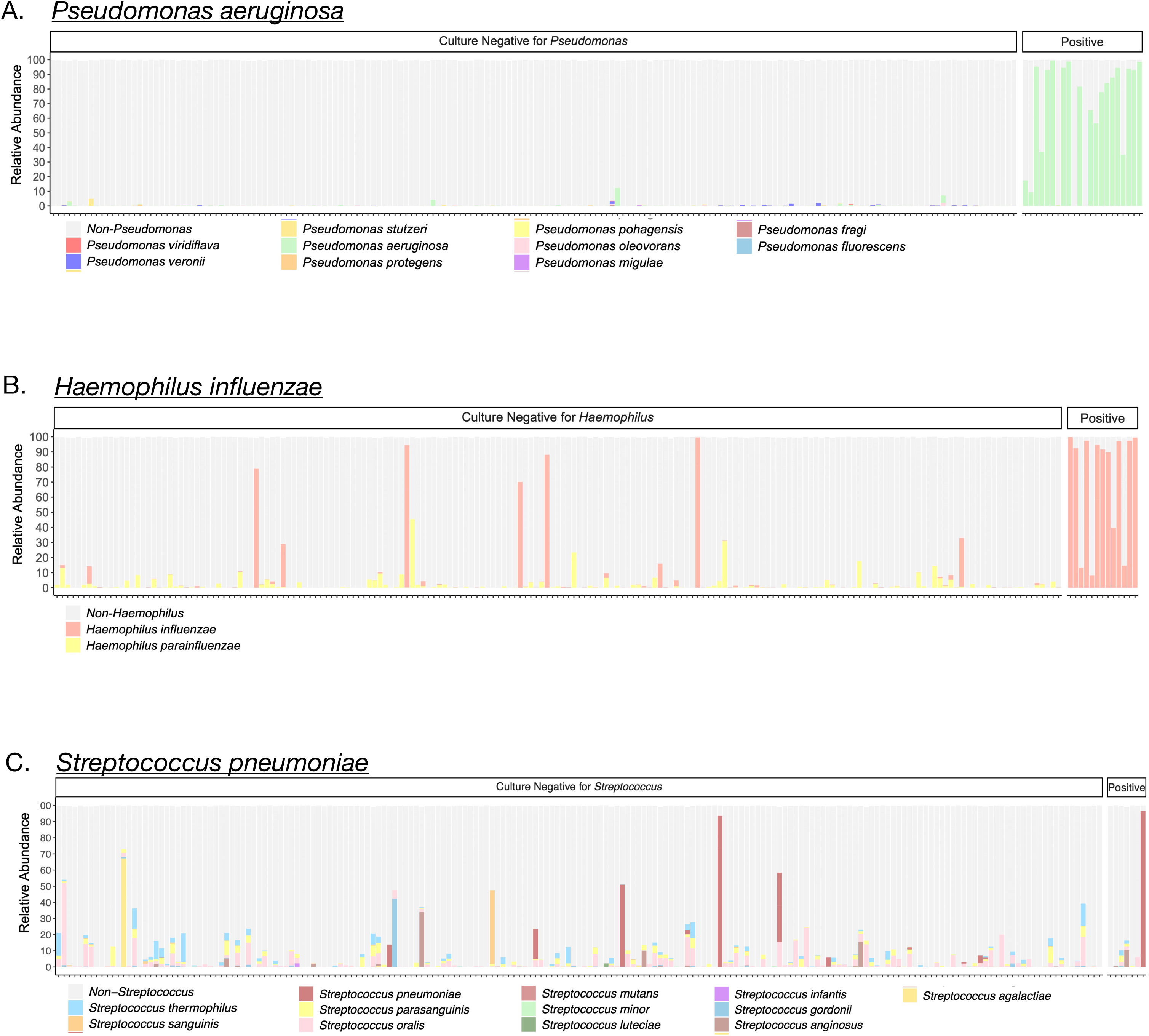
Overlap of 16S rRNA and culture for secondary pathogens. Abundance of ASVs (16SrRNA) annotated to *Pseudomonas*, *Haemophilus* and *Streptococci* in individual BAL samples showed the strongest association with culture positivity for *Pseudomonas aeruginosa* **(Fig. E5A),** followed by *Haemophilus influenzae* **(Fig. E5B),** and then *Streptococcus pneumoniae* **(Fig. E5C).**

**Figure E6.**
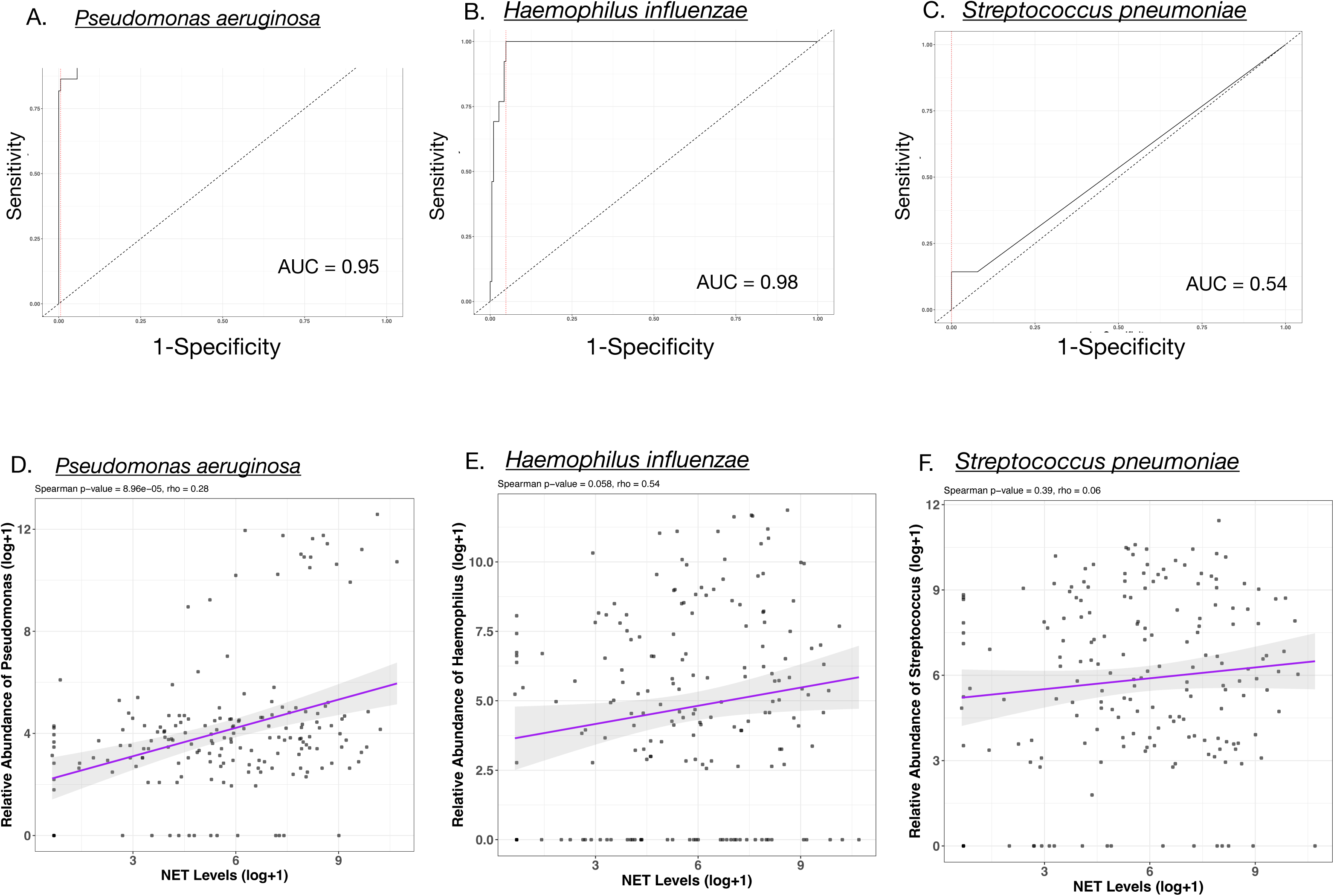
Relative abundance, culture positivity and NET levels for secondary pathogens. Receiver-operator characteristic curves comparing 16S rRNA with culture positivity showed an AUC of 0.95 and 0.98, respectively, for *Pseudomonas* and *Haemophilus* **(Fig. E6A-B)** and an AUC of 0.54 for *Streptococcus* **(Fig. E6C).** Direct comparison of the relative abundances of *Pseudomonas, Haemophilus* and *Streptococcus* with NET levels showed a highly significant Spearman’s rank correlation coefficient for *Pseudomonas* (Spearman’s p= 8.96e-05, **Fig. E6D),** but the coefficient was not significant for *Haemophilus* or *Streptococcus* **(Fig. E6E-F).**

**Figure E7.**
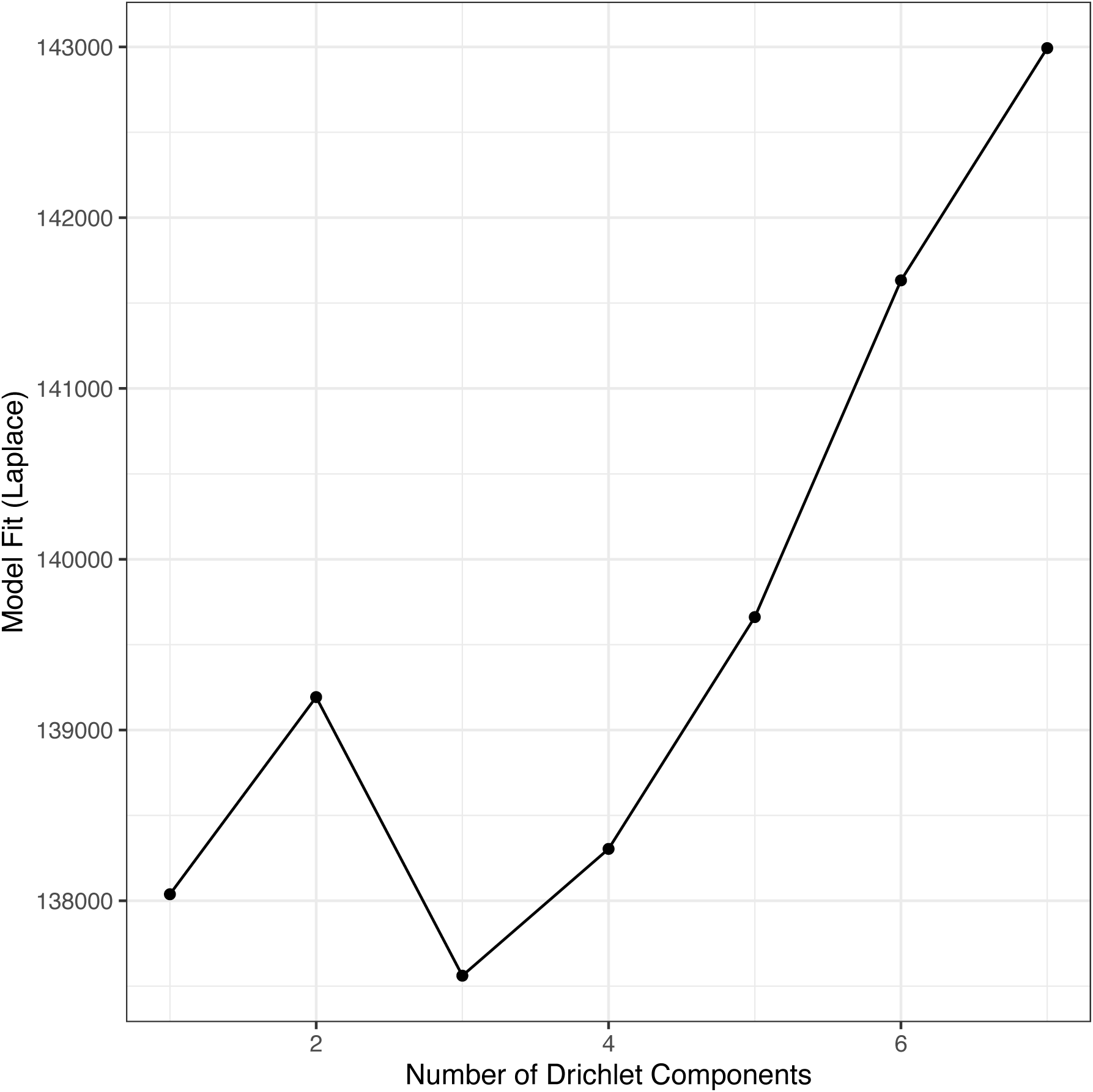
**Dirichlet Multinomial Modeling identified three possible clusters,** or microbial metacommunities in the lower airways that had the best fit for the 16S rRNA data.

**Figure E8.**
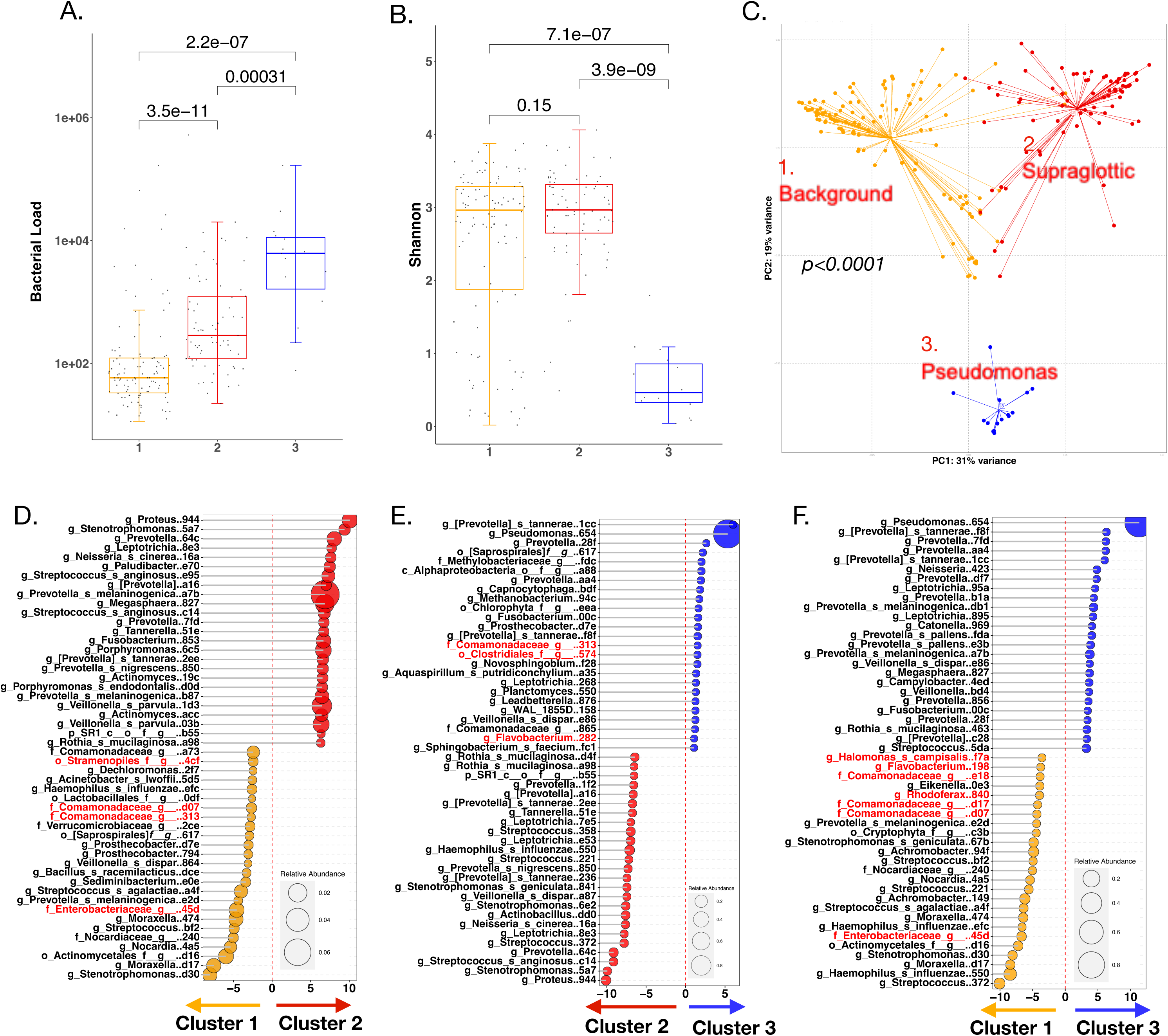
Microbial composition of lower airway DMM clusters. Bacterial load was significantly different between all three clusters (1 versus 2: p<0.0001; 2 versus 3: p=0.00031 and 1 versus 3: p<0.0001, **Fig. E8A).** Cluster 1 had the lowest bacterial load whereas cluster 3 had the highest. Alpha diversity was also significantly different between the clusters (2 versus 3: p<0.0001 and 1 versus 3: p<0.0001, **Fig. E8B**). Cluster 1 and 2 had higher alpha diversities than cluster 3. Bray Curtis analyses showed all three clusters to be compositionally distinct (p<0.0001, PERMANOVA, **Fig. E8C**). Cluster 2 was enriched with oral commensals such as *Prevotella*, *Streptococcus* and *Veillonella* (**Fig. E8D-E**) and cluster 3 was enriched with *Pseudomonas* (**Fig. E8F**).

**Figure E9.**
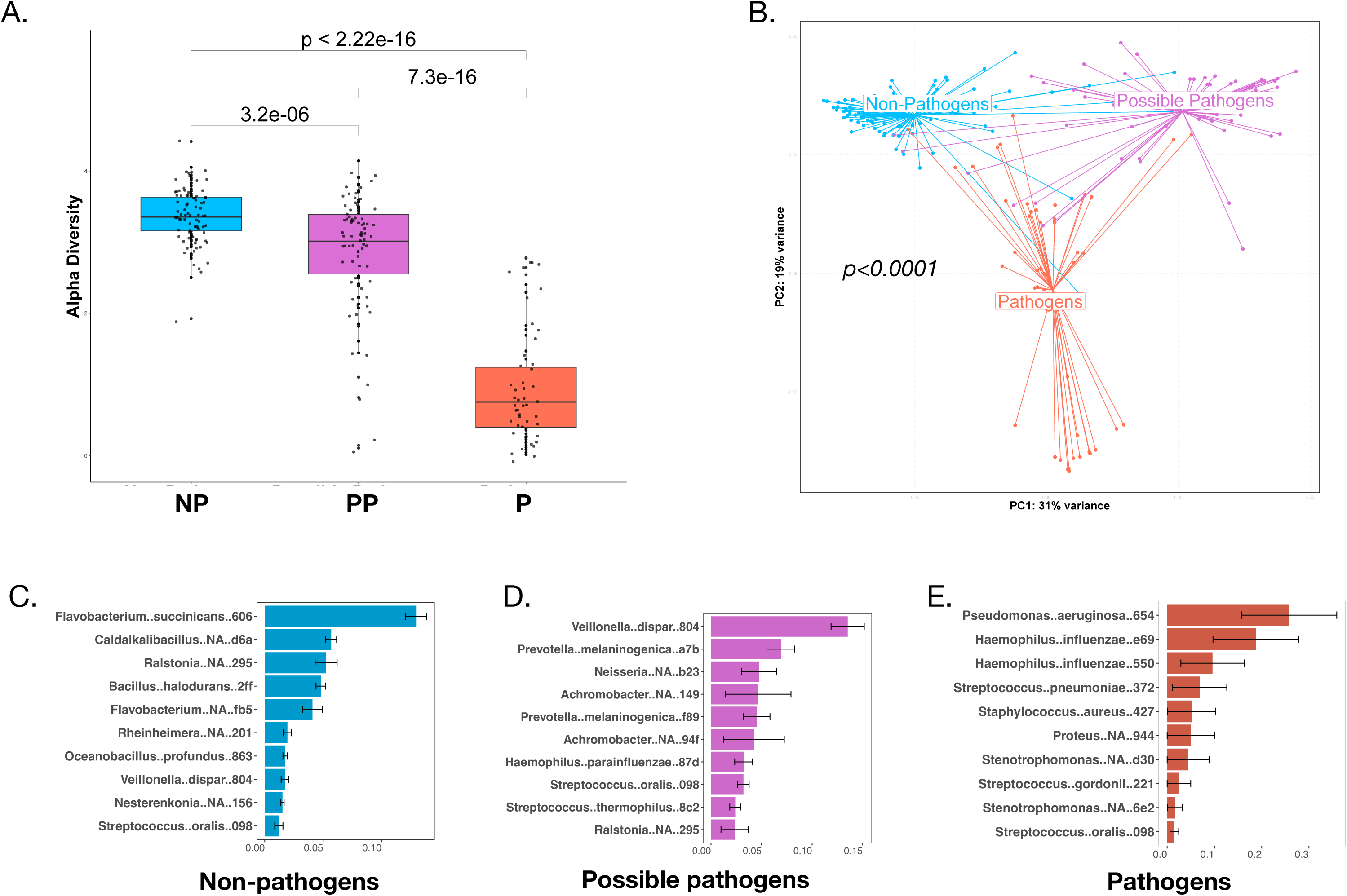
Lower airway clusters based on pathogenic potential. We categorized all 16S rRNA annotations based on pathogenic potential at the genus level as pathogens (P), possible-pathogens (PP) and non-pathogens (PP). **Fig. E9A.** Differences in alpha diversity were significant between the NP, PP and P groups; with the lowest value for the P and highest for the NP cluster (p<0.0001 for all comparison, Mann Whitney). **Fig. E9B.** Beta diversity was also significantly different between the three clusters (p<0.0001, PERMANOVA). **Fig. E6C.** The NP cluster was enriched with *Flavobacterium.* **Fig. E9D.** The PP cluster was enriched with oral commensals such as *Prevotella* and *Veillonella*. **Fig. E9E.** The P cluster and was enriched with *Pseudomonas*, *Haemophilus* and *Streptococcus*.

**Figure E10.**
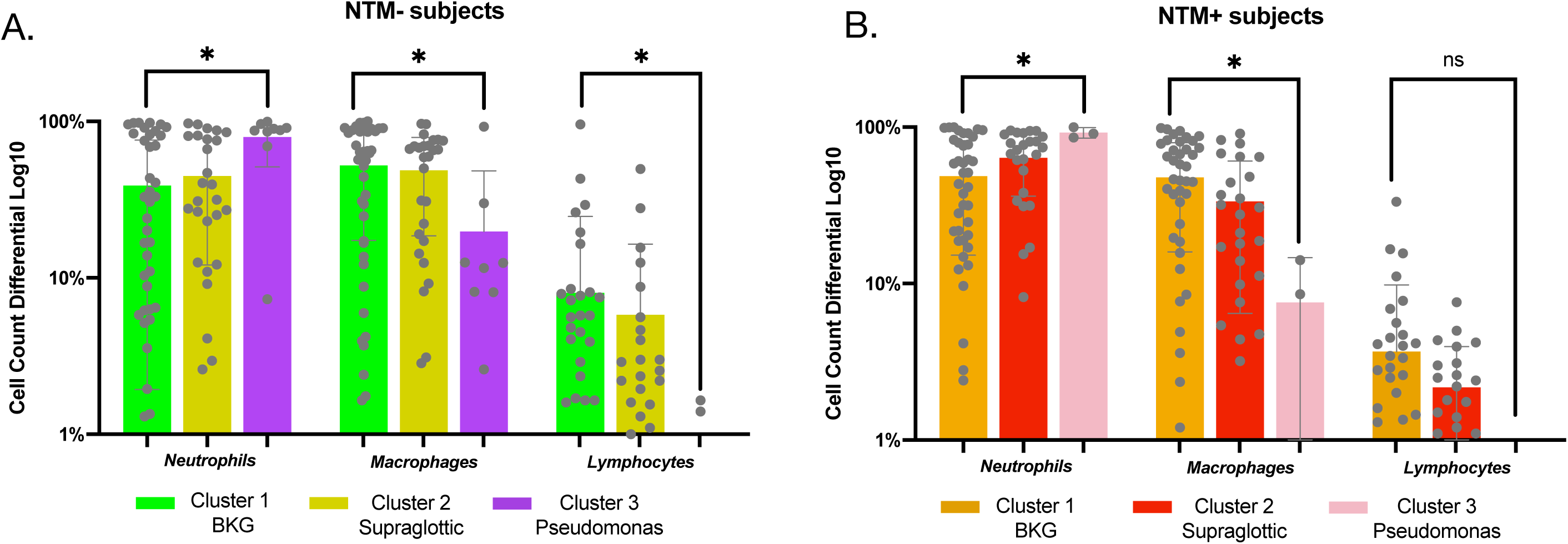
Cluster 3 group had high neutrophils. Fig. E10A-B. In both NTM- and NTM+ groups, the DMM cluster 3 had significantly higher neutrophil counts and significantly lower macrophage counts, as compared to both cluster 1 and cluster 2 (p<0.05, Mann Whitney).

**Figure E11.**
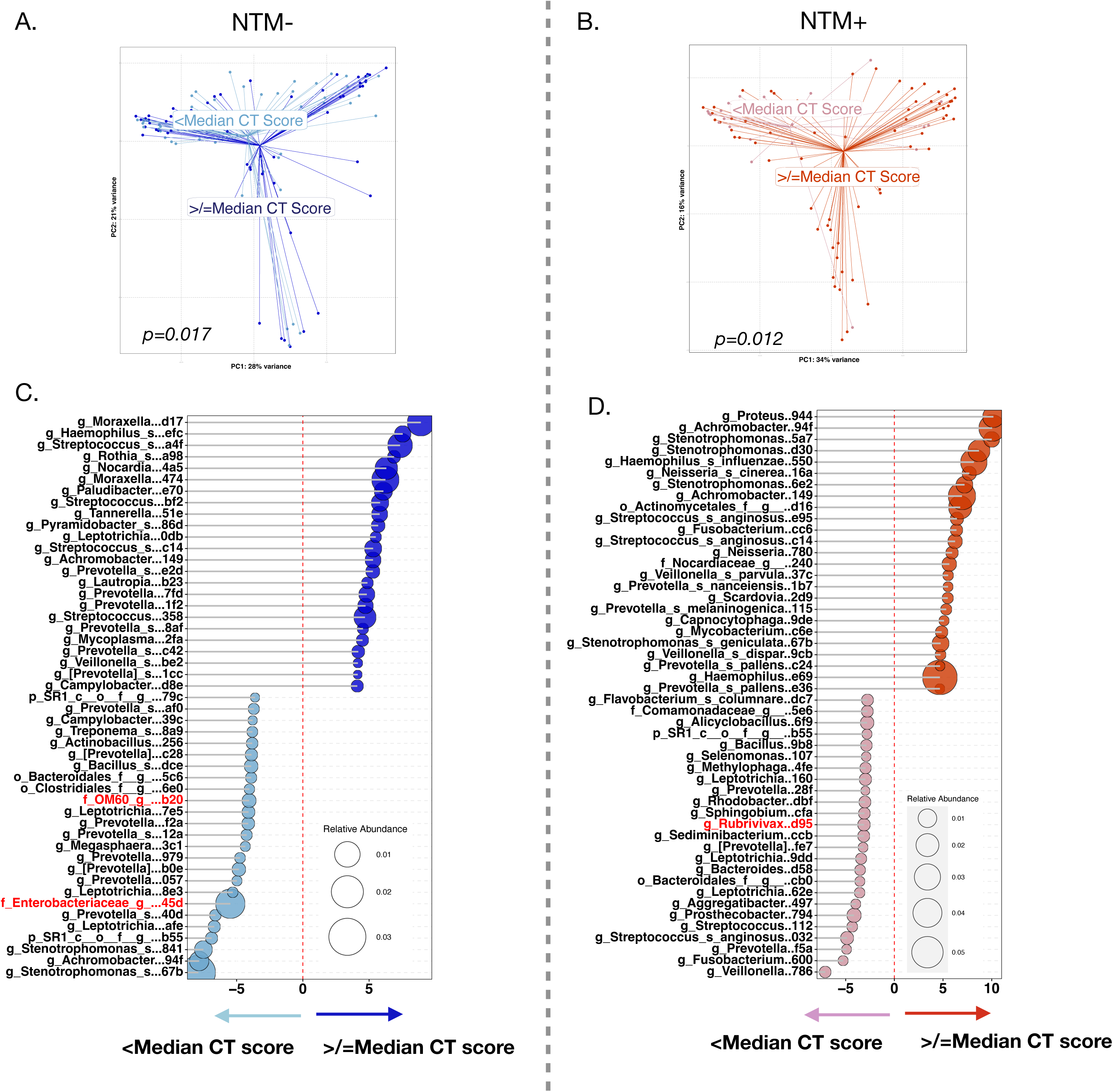
NTM+ BAL with a higher-than-median Chest CT score was more enriched with oral commensals. Fig. E11A-B. Using Chest CT scores, we found significant compositional differences between NTM-/NTM+ subjects (p=0.017 and 0.012 respectively, PERMANOVA). **Fig. E11C-D.** The relative abundance of *Haemophilus, Achromobacter* and oral commensals such as *Veillonella, Prevotella, Stenotrophomonas*, *Proteus* were significantly higher in the NTM+ “above-median-CT-score” group. The NTM-“above-median-CT-score” group showed some *Nocardia* and *Streptococcus* ASVs.

**Figure E12.**
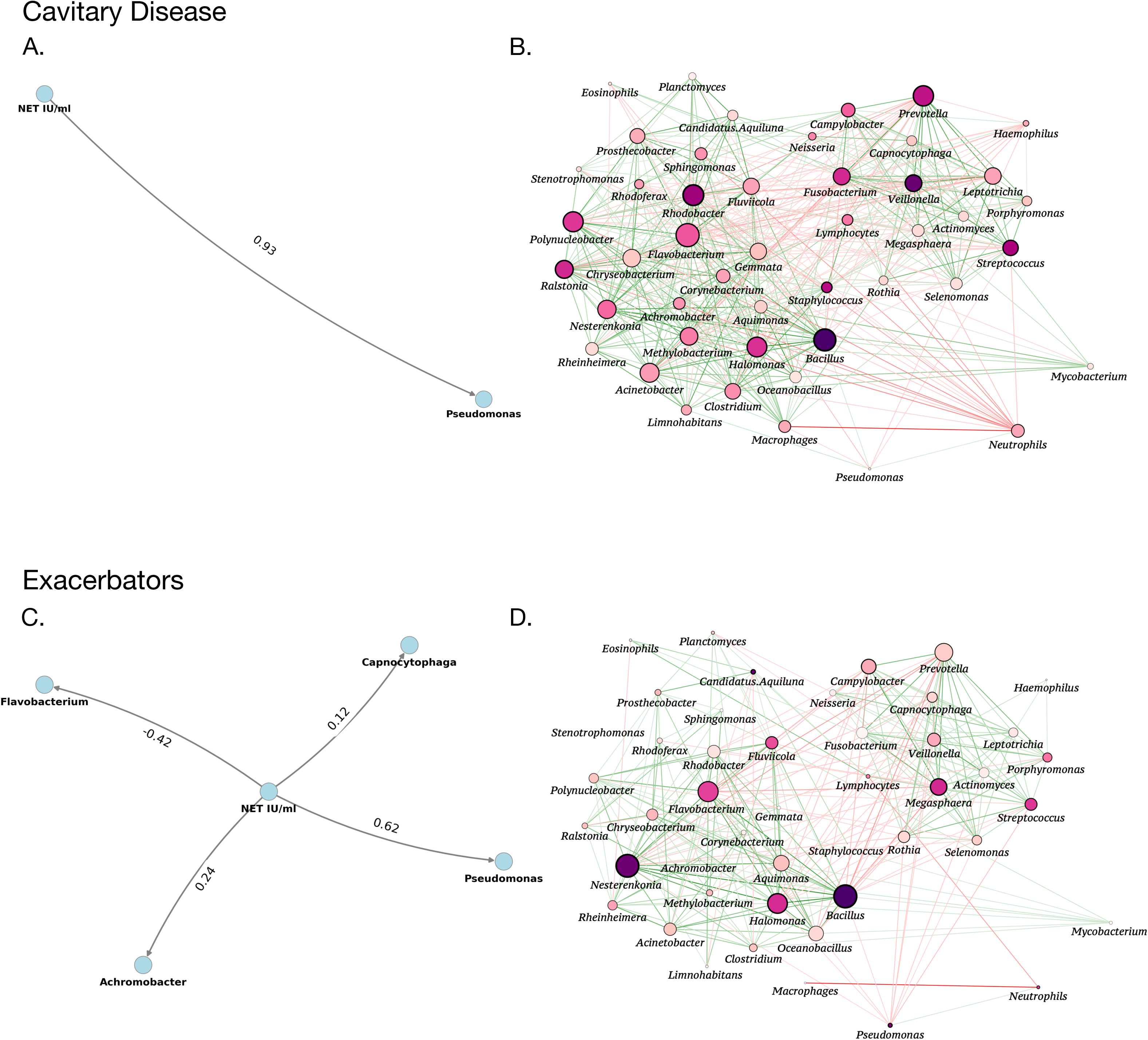
NET is associated with *Pseudomonas*, oral commensals and *Mycobacteria* in severe disease phenotypes. Fig. E12A. MaAsLin2 analysis showed NET levels had a positive interaction with *Pseudomonas*, (coefficient of association 0.93) in cavitary disease. **Fig. E12B.** *Pseudomonas* exhibited competitive interactions with oral commensals (red lines) whereas *Mycobacteria* was synchronous with several oral commensals such as *Streptococcus* and *Prevotella* (green lines). **Fig. E12C.** MaAsLin2 analysis showed that in exacerbators, NET levels had a positive interaction with *Pseudomonas* and *Achromobacter* (coefficient of association 0.62 and 0.24, respectively) and a negative one with *Flavobacterium* (coefficient of association -0.42). **Fig. E12D.** *Pseudomonas* had a competitive interaction with oral commensals (red lines) whereas *Mycobacteria* was synchronous with several oral commensals (green lines).

**Figure E13.**
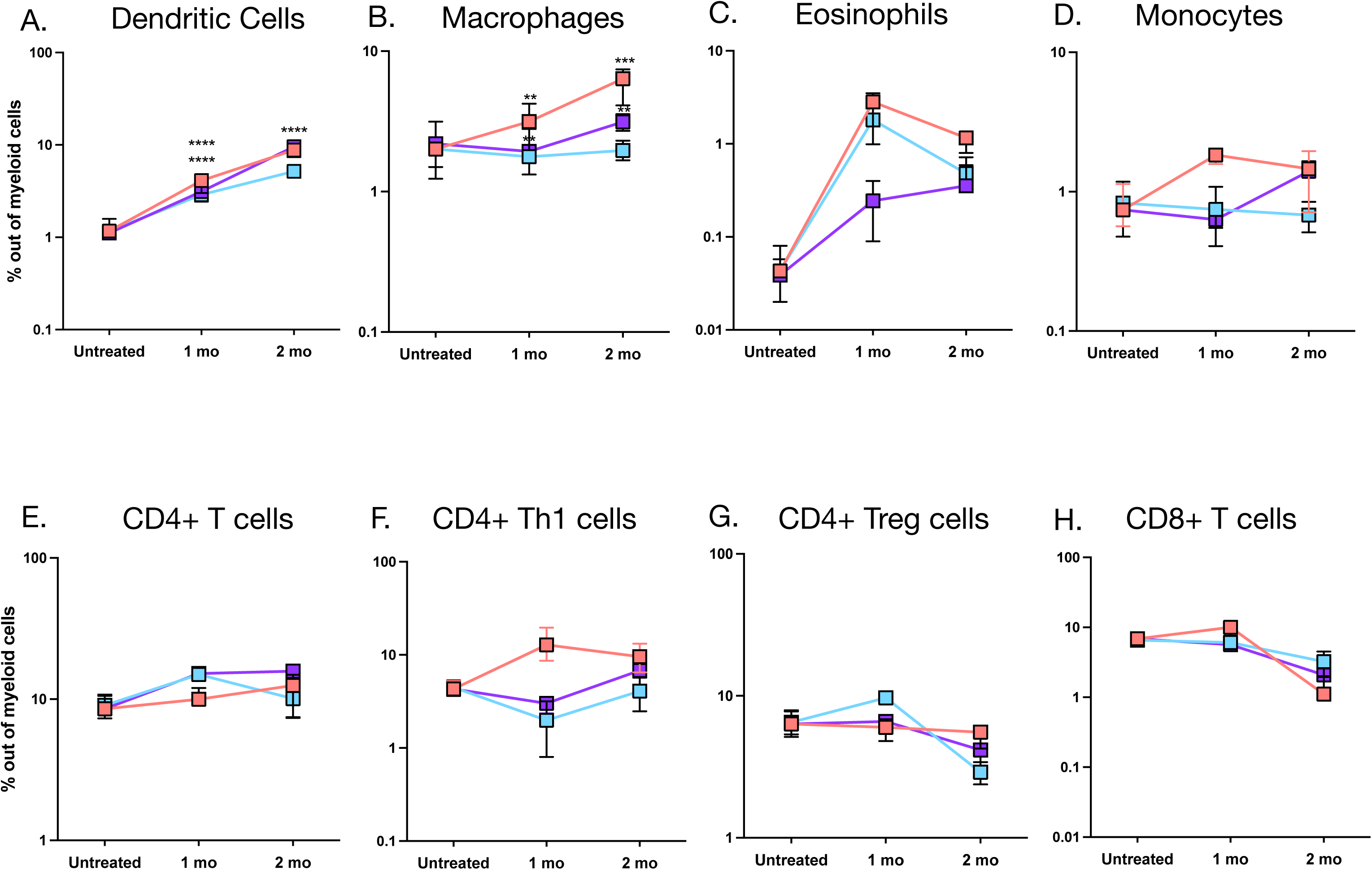
NTM upregulates dendritic cells and macrophages in a murine micro-aspiration model. Fig. E13A. NTM infection drove the influx of dendritic cells 1 month (p<0.0001, One-way ANOVA) and 2 months (p<0.0001, One-way ANOVA). This NTM-driven influx was augmented by MOC at 1 month (p<0.0001, One-way ANOVA). **Fig. E13B.** NTM infection drove the influx of macrophages at 1 month (p=0.0012, One-way ANOVA) and 2 months (p=0.0003, One-way ANOVA). This NTM-driven influx was augmented by MOC at both at 1 and 2 months (p=0.0022 and p=0.0047, respectively, One-way ANOVA). **Fig. E13C-F.** No significant trends were noted at either time points for eosinophils, monocytes, CD4+ T cells or Th1 cells. **Fig. E13G-H.** Both T_REG_ cells and CD8+ T cells seemed to down-trend, but this was not significant.

**Figure E14.**
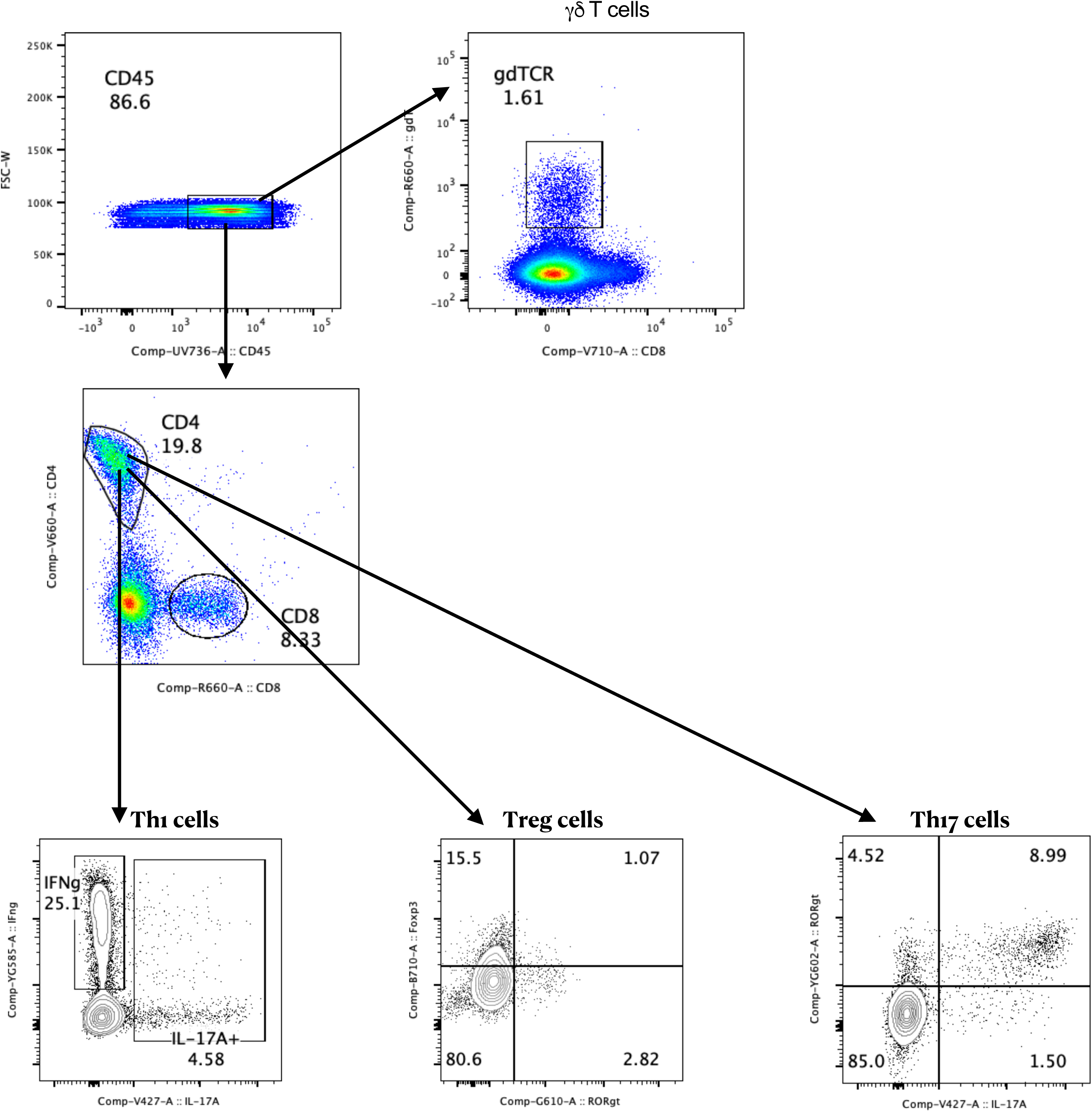
Gating strategy for T cell subsets. After exclusion of doublets using a combination of side scatter height (SSC-H) and side scatter area (SSC-A), forward scatter height (FSC-H) and forward scatter width (FSC-W), were used to gate for CD45.2+ cells. This selected total leukocytes. For T cell subsets, CD4+ T-cells, CD8+ T-cells, and γ8TCR+ T-cells were identified based on the expression of CD4, CD8, and γ8TCR, respectively. CD4+Th1, CD4+T_REG_, and CD4+Th17 cells were identified based on expression CD4+IFN-γ+, CD4+Foxp3+, CD4+IL-17A+RORγt+ cells respectively. CD8+ effector cells were identified based on CD8+IFN-γ+ cells.

**Figure E15.**
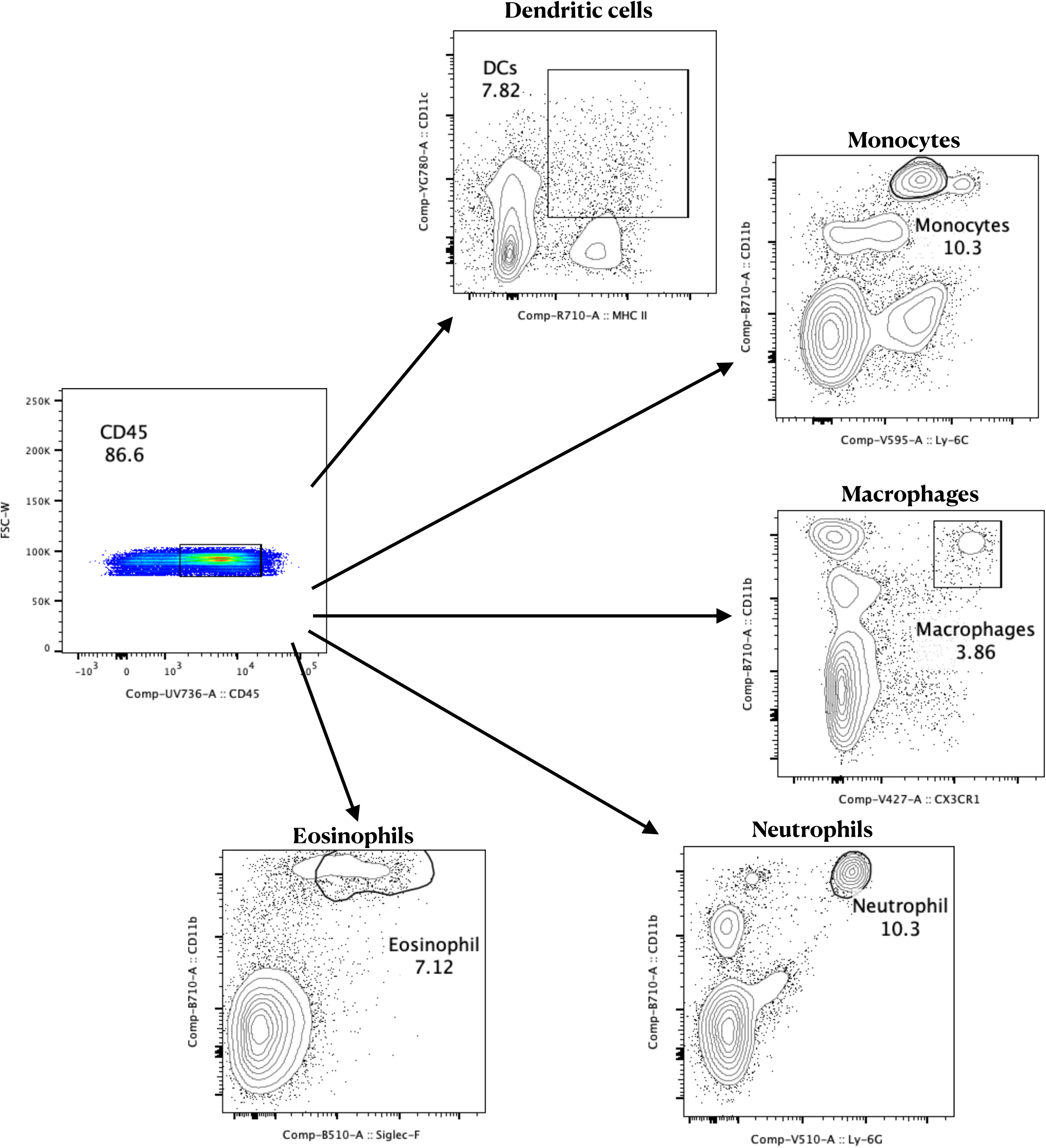
Gating strategy for myeloid subsets. For myeloid subsets within CD45.2+CD11b+ cells neutrophils, macrophage, monocytes and eosinophils were identified as Ly-6G^+^, CX3CR1^+^, Ly-6C^+^, or Siglec-F^+^ cells, respectively. Dendritic cells were identified as CD45.2^+^CD11c^+^MHC II^+^ cells

## References

1. **Boyton RJ, Altmann DM. Bronchiectasis: Current Concepts in Pathogenesis, Immunology, and Microbiology. Annu Rev Pathol 2016; 11: 523–554.

2. Diagnosis and treatment of disease caused by nontuberculous mycobacteria. This official statement of the American Thoracic Society was approved by the Board of Directors, March 1997. Medical Section of the American Lung Association. Am J Respir Crit Care Med 1997; 156: S1–25.

3. Zhou Y, Mu W, Zhang J, Wen SW, Pakhale S. Global prevalence of non-tuberculous mycobacteria in adults with non-cystic fibrosis bronchiectasis 2006-2021: a systematic review and meta-analysis. BMJ Open 2022; 12: e055672.

4. Strollo SE, Adjemian J, Adjemian MK, Prevots DR. The Burden of Pulmonary Nontuberculous Mycobacterial Disease in the United States. Ann Am Thorac Soc 2015; 12: 1458–1464.

5. Brinkmann V, Reichard U, Goosmann C, Fauler B, Uhlemann Y, Weiss DS, Weinrauch Y, Zychlinsky A. Neutrophil extracellular traps kill bacteria. Science (New York, NY) 2004; 303: 1532–1535.

6. Keir HR, Shoemark A, Dicker AJ, Perea L, Pollock J, Giam YH, Suarez-Cuartin G, Crichton ML, Lonergan M, Oriano M, Cant E, Einarsson GG, Furrie E, Elborn JS, Fong CJ, Finch S, Rogers GB, Blasi F, Sibila O, Aliberti S, Simpson JL, Huang JTJ, Chalmers JD. Neutrophil extracellular traps, disease severity, and antibiotic response in bronchiectasis: an international, observational, multicohort study. The Lancet Respiratory medicine 2021; 9: 873–884.

7. Chalmers JD, Haworth CS, Metersky ML, Loebinger MR, Blasi F, Sibila O, O’Donnell AE, Sullivan EJ, Mange KC, Fernandez C, Zou J, Daley CL, Investigators W. Phase 2 Trial of the DPP-1 Inhibitor Brensocatib in Bronchiectasis. N Engl J Med 2020; 383: 2127–2137.

8. Chalmers JD, Gupta A, Chotirmall SH, Armstrong A, Eickholz P, Hasegawa N, McShane PJ, O’Donnell AE, Shteinberg M, Watz H, Eleftheraki A, Diefenbach C, Sauter W. A Phase 2 randomised study to establish efficacy, safety and dosing of a novel oral cathepsin C inhibitor, BI 1291583, in adults with bronchiectasis: Airleaf. ERJ Open Res 2023; 9.

9. Chalmers JD, Burgel PR, Daley CL, De Soyza A, Haworth CS, Mauger D, Mange K, Teper A, Fernandez C, Conroy D, Metersky M. Brensocatib in non-cystic fibrosis bronchiectasis: ASPEN protocol and baseline characteristics. ERJ Open Res 2024; 10.

10. Chalmers JD, Burgel PR, Daley CL, De Soyza A, Haworth CS, Mauger D, Loebinger MR, McShane PJ, Ringshausen FC, Blasi F, Shteinberg M, Mange K, Teper A, Fernandez C, Zambrano M, Fan C, Zhang X, Metersky ML. Phase 3 Trial of the DPP-1 Inhibitor Brensocatib in Bronchiectasis. The New England journal of medicine 2025; 392: 1569–1581.

11. Nakamura K, Nakayama H, Sasaki S, Takahashi K, Iwabuchi K. Mycobacterium avium-intracellulare complex promote release of pro-inflammatory enzymes matrix metalloproteinases by inducing neutrophil extracellular trap formation. Scientific reports 2022; 12: 5181.

12. *FACED Martínez-García M, de Gracia J, Vendrell Relat M, Girón RM, Máiz Carro L, de la Rosa Carrillo D, Olveira C. Multidimensional approach to non-cystic fibrosis bronchiectasis: the FACED score. Eur Respir J 2014; 43: 1357–1367.

13. Chalmers JD, Aliberti S, Filonenko A, Shteinberg M, Goeminne PC, Hill AT, Fardon TC, Obradovic D, Gerlinger C, Sotgiu G, Operschall E, Rutherford RM, Dimakou K, Polverino E, De Soyza A, McDonnell MJ. Characterization of the "Frequent Exacerbator Phenotype" in Bronchiectasis. American journal of respiratory and critical care medicine 2018; 197: 1410–1420.

14. Polverino E, Goeminne PC, McDonnell MJ, Aliberti S, Marshall SE, Loebinger MR, Murris M, Cantón R, Torres A, Dimakou K, De Soyza A, Hill AT, Haworth CS, Vendrell M, Ringshausen FC, Subotic D, Wilson R, Vilaró J, Stallberg B, Welte T, Rohde G, Blasi F, Elborn S, Almagro M, Timothy A, Ruddy T, Tonia T, Rigau D, Chalmers JD. European Respiratory Society guidelines for the management of adult bronchiectasis. Eur Respir J 2017; 50.

15. Daley CL, Iaccarino JM, Lange C, Cambau E, Wallace RJ, Jr., Andrejak C, Böttger EC, Brozek J, Griffith DE, Guglielmetti L, Huitt GA, Knight SL, Leitman P, Marras TK, Olivier KN, Santin M, Stout JE, Tortoli E, van Ingen J, Wagner D, Winthrop KL. Treatment of nontuberculous mycobacterial pulmonary disease: an official ATS/ERS/ESCMID/IDSA clinical practice guideline. Eur Respir J 2020; 56.

16. Robinson MD, McCarthy DJ, Smyth GK. edgeR: a Bioconductor package for differential expression analysis of digital gene expression data. Bioinformatics 2010; 26: 139–140.

17. McCarthy DJ, Chen Y, Smyth GK. Differential expression analysis of multifactor RNA-Seq experiments with respect to biological variation. Nucleic Acids Res 2012; 40: 4288–4297.

18. Chen Y, Lun AT, Smyth GK. From reads to genes to pathways: differential expression analysis of RNA-Seq experiments using Rsubread and the edgeR quasi-likelihood pipeline. F1000Res 2016; 5: 1438.

19. Holmes I, Harris K, Quince C. Dirichlet multinomial mixtures: generative models for microbial metagenomics. PloS one 2012; 7: e30126.

20. Mac Aogain M, Narayana JK, Tiew PY, Ali N, Yong VFL, Jaggi TK, Lim AYH, Keir HR, Dicker AJ, Thng KX, Tsang A, Ivan FX, Poh ME, Oriano M, Aliberti S, Blasi F, Low TB, Ong TH, Oliver B, Giam YH, Tee A, Koh MS, Abisheganaden JA, Tsaneva-Atanasova K, Chalmers JD, Chotirmall SH. Integrative microbiomics in bronchiectasis exacerbations. Nat Med 2021; 27: 688–699.

21. Narayana JK, Aliberti S, Mac Aogain M, Jaggi TK, Ali N, Ivan FX, Cheng HS, Yip YS, Vos MIG, Low ZS, Lee JXT, Amati F, Gramegna A, Wong SH, Sung JJY, Tan NS, Tsaneva-Atanasova K, Blasi F, Chotirmall SH. Microbial Dysregulation of the Gut-Lung Axis in Bronchiectasis. Am J Respir Crit Care Med 2023; 207: 908–920.

22. Mallick H, Rahnavard A, McIver LJ, Ma S, Zhang Y, Nguyen LH, Tickle TL, Weingart G, Ren B, Schwager EH, Chatterjee S, Thompson KN, Wilkinson JE, Subramanian A, Lu Y, Waldron L, Paulson JN, Franzosa EA, Bravo HC, Huttenhower C. Multivariable association discovery in population-scale meta-omics studies. PLoS Comput Biol 2021; 17: e1009442.

23. Wu BG, Sulaiman I, Tsay JJ, Perez L, Franca B, Li Y, Wang J, Gonzalez AN, El-Ashmawy M, Carpenito J, Olsen E, Sauthoff M, Yie K, Liu X, Shen N, Clemente JC, Kapoor B, Zangari T, Mezzano V, Loomis C, Weiden MD, Koralov SB, D’Armiento J, Ahuja SK, Wu XR, Weiser JN, Segal LN. Episodic Aspiration with Oral Commensals Induces a MyD88-dependent, Pulmonary T-Helper Cell Type 17 Response that Mitigates Susceptibility to Streptococcus pneumoniae. American journal of respiratory and critical care medicine 2021; 203: 1099–1111.

24. Tsay JJ, Wu BG, Sulaiman I, Gershner K, Schluger R, Li Y, Yie TA, Meyn P, Olsen E, Perez L, Franca B, Carpenito J, Iizumi T, El-Ashmawy M, Badri M, Morton JT, Shen N, He L, Michaud G, Rafeq S, Bessich JL, Smith RL, Sauthoff H, Felner K, Pillai R, Zavitsanou AM, Koralov SB, Mezzano V, Loomis CA, Moreira AL, Moore W, Tsirigos A, Heguy A, Rom WN, Sterman DH, Pass HI, Clemente JC, Li H, Bonneau R, Wong KK, Papagiannakopoulos T, Segal LN. Lower Airway Dysbiosis Affects Lung Cancer Progression. Cancer Discov 2021; 11: 293–307.

25. Gautam S, Stahl Y, Young GM, Howell R, Cohen AJ, Tsang DA, Martin T, Sharma L, Dela Cruz CS. Quantification of bronchoalveolar neutrophil extracellular traps and phagocytosis in murine pneumonia. American journal of physiology Lung cellular and molecular physiology 2020; 319: L661–l669.

26. Dicker AJ, Crichton ML, Pumphrey EG, Cassidy AJ, Suarez-Cuartin G, Sibila O, Furrie E, Fong CJ, Ibrahim W, Brady G, Einarsson GG, Elborn JS, Schembri S, Marshall SE, Palmer CNA, Chalmers JD. Neutrophil extracellular traps are associated with disease severity and microbiota diversity in patients with chronic obstructive pulmonary disease. The Journal of allergy and clinical immunology 2018; 141: 117–127.

27. Sulaiman I, Wu BG, Li Y, Scott AS, Malecha P, Scaglione B, Wang J, Basavaraj A, Chung S, Bantis K, Carpenito J, Clemente JC, Shen N, Bessich J, Rafeq S, Michaud G, Donington J, Naidoo C, Theron G, Schattner G, Garofano S, Condos R, Kamelhar D, Addrizzo-Harris D, Segal LN. Evaluation of the airway microbiome in nontuberculous mycobacteria disease. Eur Respir J 2018; 52.

28. Claeys TA, Rosas Mejia O, Marshall S, Jarzembowski JA, Hayes D, Jr., Hull NM, Liyanage NPM, Chun RH, Sulman CG, Huppler AR, Robinson RT. Attenuation of Helper T Cell Capacity for TH1 and TH17 Differentiation in Children With Nontuberculous Mycobacterial Infection. J Infect Dis 2019; 220: 1843–1847.

29. Schuurbiers MMF, Bruno M, Zweijpfenning SMH, Magis-Escurra C, Boeree M, Netea MG, van Ingen J, van de Veerdonk F, Hoefsloot W. Immune defects in patients with pulmonary Mycobacterium abscessus disease without cystic fibrosis. ERJ Open Res 2020; 6.

30. Vázquez N, Rekka S, Gliozzi M, Feng CG, Amarnath S, Orenstein JM, Wahl SM. Modulation of innate host factors by Mycobacterium avium complex in human macrophages includes interleukin 17. J Infect Dis 2012; 206: 1206–1217.

31. Zhang Y, Chandra V, Riquelme Sanchez E, Dutta P, Quesada PR, Rakoski A, Zoltan M, Arora N, Baydogan S, Horne W, Burks J, Xu H, Hussain P, Wang H, Gupta S, Maitra A, Bailey JM, Moghaddam SJ, Banerjee S, Sahin I, Bhattacharya P, McAllister F. Interleukin-17-induced neutrophil extracellular traps mediate resistance to checkpoint blockade in pancreatic cancer. The Journal of experimental medicine 2020; 217.

32. de Boer OJ, Li X, Teeling P, Mackaay C, Ploegmakers HJ, van der Loos CM, Daemen MJ, de Winter RJ, van der Wal AC. Neutrophils, neutrophil extracellular traps and interleukin-17 associate with the organisation of thrombi in acute myocardial infarction. Thromb Haemost 2013; 109: 290–297.

33. Papagoras C, Chrysanthopoulou A, Mitsios A, Ntinopoulou M, Tsironidou V, Batsali AK, Papadaki HA, Skendros P, Ritis K. IL-17A expressed on neutrophil extracellular traps promotes mesenchymal stem cell differentiation toward bone-forming cells in ankylosing spondylitis. European journal of immunology 2021; 51: 930–942.

34. Lindén A, Laan M, Anderson GP. Neutrophils, interleukin-17A and lung disease. Eur Respir J 2005; 25: 159–172.

35. Kang I, Kim Y, Lee HK. Double-edged sword: γδ T cells in mucosal homeostasis and disease. Exp Mol Med 2023; 55: 1895–1904.

36. Shu CC, Pan SW, Feng JY, Wang JY, Chan YJ, Yu CJ, Su WJ. The Clinical Significance of Programmed Death-1, Regulatory T Cells and Myeloid Derived Suppressor Cells in Patients with Nontuberculous Mycobacteria-Lung Disease. Journal of clinical medicine 2019; 8.

37. Dicker AJ, Lonergan M, Keir HR, Smith AH, Pollock J, Finch S, Cassidy AJ, Huang JTJ, Chalmers JD. The sputum microbiome and clinical outcomes in patients with bronchiectasis: a prospective observational study. The Lancet Respiratory medicine 2021; 9: 885–896.

38. Araújo D, Shteinberg M, Aliberti S, Goeminne PC, Hill AT, Fardon TC, Obradovic D, Stone G, Trautmann M, Davis A, Dimakou K, Polverino E, De Soyza A, McDonnell MJ, Chalmers JD. The independent contribution of Pseudomonas aeruginosa infection to long-term clinical outcomes in bronchiectasis. Eur Respir J 2018; 51.

39. Narayana JK, Mac Aogáin M, Ali N, Tsaneva-Atanasova K, Chotirmall SH. Similarity network fusion for the integration of multi-omics and microbiomes in respiratory disease. Eur Respir J 2021; 58.

40. Segal LN, Clemente JC, Tsay J-CJ, Koralov SB, Keller BC, Wu BG, Li Y, Shen N, Ghedin E, Morris A, Diaz P, Huang L, Wikoff WR, Ubeda C, Artacho A, Rom WN, Sterman DH, Collman RG, Blaser MJ, Weiden MD. Enrichment of the lung microbiome with oral taxa is associated with lung inflammation of a Th17 phenotype. Nature Microbiology 2016: 16031.

41. Segal LN, Alekseyenko AV, Clemente JC, Kulkarni R, Wu B, Gao Z, Chen H, Berger KI, Goldring RM, Rom WN, Blaser MJ, Weiden MD. Enrichment of lung microbiome with supraglottic taxa is associated with increased pulmonary inflammation. Microbiome 2013; 1: 19.

42. Segal LN, Alekseyenko AV, Clemente JC, Kulkarni R, Wu B, Chen H, Berger KI, Goldring RM, Rom WN, Blaser MJ, Weiden MD. Enrichment of lung microbiome with supraglottic taxa is associated with increased pulmonary inflammation. Microbiome 2013; 1: 19.

43. Li L, Mac Aogáin M, Xu T, Jaggi TK, Chan LLY, Qu J, Wei L, Liao S, Cheng HS, Keir HR, Dicker AJ, Tan KS, De Yun W, Koh MS, Ong TH, Lim AYH, Abisheganaden JA, Low TB, Hassan TM, Long X, Wark PAB, Oliver B, Drautz-Moses DI, Schuster SC, Tan NS, Fang M, Chalmers JD, Chotirmall SH. Neisseria species as pathobionts in bronchiectasis. Cell host & microbe 2022; 30: 1311–1327.e1318.

44. Singh S, Segal LN. A lung pathobiont story: Thinking outside the Koch’s postulate box. Cell host & microbe 2022; 30: 1196–1198.

45. Konstan MW, Döring G, Heltshe SL, Lands LC, Hilliard KA, Koker P, Bhattacharya S, Staab A, Hamilton A. A randomized double blind, placebo controlled phase 2 trial of BIIL 284 BS (an LTB4 receptor antagonist) for the treatment of lung disease in children and adults with cystic fibrosis. J Cyst Fibros 2014; 13: 148–155.

46. De Soyza A, Pavord I, Elborn JS, Smith D, Wray H, Puu M, Larsson B, Stockley R. A randomised, placebo-controlled study of the CXCR2 antagonist AZD5069 in bronchiectasis. Eur Respir J 2015; 46: 1021–1032.

47. **Verma D, Stapleton M, Gadwa J, Vongtongsalee K, Schenkel AR, Chan ED, Ordway D. Mycobacterium avium Infection in a C3HeB/FeJ Mouse Model. Front Microbiol 2019; 10: 693.

48. Tunney MM, Einarsson GG, Wei L, Drain M, Klem ER, Cardwell C, Ennis M, Boucher RC, Wolfgang MC, Elborn JS. Lung microbiota and bacterial abundance in patients with bronchiectasis when clinically stable and during exacerbation. American journal of respiratory and critical care medicine 2013; 187: 1118–1126.

49. Byun MK, Chang J, Kim HJ, Jeong SH. Differences of lung microbiome in patients with clinically stable and exacerbated bronchiectasis. PloS one 2017; 12: e0183553.

50. López-Roa P, Esteban J, Muñoz-Egea MC. Updated Review on the Mechanisms of Pathogenicity in Mycobacterium abscessus, a Rapidly Growing Emerging Pathogen. Microorganisms 2022; 11.

51. Malcolm KC, Caceres SM, Pohl K, Poch KR, Bernut A, Kremer L, Bratton DL, Herrmann JL, Nick JA. Neutrophil killing of Mycobacterium abscessus by intra- and extracellular mechanisms. PloS one 2018; 13: e0196120.

52. To K, Cao R, Yegiazaryan A, Owens J, Venketaraman V. General Overview of Nontuberculous Mycobacteria Opportunistic Pathogens: Mycobacterium avium and Mycobacterium abscessus. Journal of clinical medicine 2020; 9.

## References

1. *FACED Martínez-García M, de Gracia J, Vendrell Relat M, Girón RM, Máiz Carro L, de la Rosa Carrillo D, Olveira C. Multidimensional approach to non-cystic fibrosis bronchiectasis: the FACED score. Eur Respir J 2014; 43: 1357–1367.

2. Chalmers JD, Aliberti S, Filonenko A, Shteinberg M, Goeminne PC, Hill AT, Fardon TC, Obradovic D, Gerlinger C, Sotgiu G, Operschall E, Rutherford RM, Dimakou K, Polverino E, De Soyza A, McDonnell MJ. Characterization of the "Frequent Exacerbator Phenotype" in Bronchiectasis. American journal of respiratory and critical care medicine 2018; 197: 1410–1420.

3. Polverino E, Goeminne PC, McDonnell MJ, Aliberti S, Marshall SE, Loebinger MR, Murris M, Cantón R, Torres A, Dimakou K, De Soyza A, Hill AT, Haworth CS, Vendrell M, Ringshausen FC, Subotic D, Wilson R, Vilaró J, Stallberg B, Welte T, Rohde G, Blasi F, Elborn S, Almagro M, Timothy A, Ruddy T, Tonia T, Rigau D, Chalmers JD. European Respiratory Society guidelines for the management of adult bronchiectasis. Eur Respir J 2017; 50.

4. Bhalla M, Turcios N, Aponte V, Jenkins M, Leitman BS, McCauley DI, Naidich DP. Cystic fibrosis: scoring system with thin-section CT. Radiology 1991; 179: 783–788.

5. Reiff DB, Wells AU, Carr DH, Cole PJ, Hansell DM. CT findings in bronchiectasis: limited value in distinguishing between idiopathic and specific types. AJR Am J Roentgenol 1995; 165: 261–267.

6. Pasteur MC, Helliwell SM, Houghton SJ, Webb SC, Foweraker JE, Coulden RA, Flower CD, Bilton D, Keogan MT. An investigation into causative factors in patients with bronchiectasis. American journal of respiratory and critical care medicine 2000; 162: 1277–1284.

7. Chalmers JD, Goeminne P, Aliberti S, McDonnell MJ, Lonni S, Davidson J, Poppelwell L, Salih W, Pesci A, Dupont LJ, Fardon TC, De Soyza A, Hill AT. The bronchiectasis severity index. An international derivation and validation study. American journal of respiratory and critical care medicine 2014; 189: 576–585.

8. Bedi P, Chalmers JD, Goeminne PC, Mai C, Saravanamuthu P, Velu PP, Cartlidge MK, Loebinger MR, Jacob J, Kamal F, Schembri N, Aliberti S, Hill U, Harrison M, Johnson C, Screaton N, Haworth C, Polverino E, Rosales E, Torres A, Benegas MN, Rossi AG, Patel D, Hill AT. The BRICS (Bronchiectasis Radiologically Indexed CT Score): A Multicenter Study Score for Use in Idiopathic and Postinfective Bronchiectasis. Chest 2018; 153: 1177–1186.

9. Helbich TH, Heinz-Peer G, Fleischmann D, Wojnarowski C, Wunderbaldinger P, Huber S, Eichler I, Herold CJ. Evolution of CT findings in patients with cystic fibrosis. AJR Am J Roentgenol 1999; 173: 81–88.

10. Song JW, Koh WJ, Lee KS, Lee JY, Chung MJ, Kim TS, Kwon OJ. High-resolution CT findings of Mycobacterium avium-intracellulare complex pulmonary disease: correlation with pulmonary function test results. AJR Am J Roentgenol 2008; 191: W160.

11. Daley CL, Iaccarino JM, Lange C, Cambau E, Wallace RJ, Jr., Andrejak C, Böttger EC, Brozek J, Griffith DE, Guglielmetti L, Huitt GA, Knight SL, Leitman P, Marras TK, Olivier KN, Santin M, Stout JE, Tortoli E, van Ingen J, Wagner D, Winthrop KL. Treatment of nontuberculous mycobacterial pulmonary disease: an official ATS/ERS/ESCMID/IDSA clinical practice guideline. Eur Respir J 2020; 56.

12. Taylor SC, Laperriere G, Germain H. Droplet Digital PCR versus qPCR for gene expression analysis with low abundant targets: from variable nonsense to publication quality data. Sci Rep 2017; 7: 2409.

13. Segal LN, Clemente JC, Tsay JC, Koralov SB, Keller BC, Wu BG, Li Y, Shen N, Ghedin E, Morris A, Diaz P, Huang L, Wikoff WR, Ubeda C, Artacho A, Rom WN, Sterman DH, Collman RG, Blaser MJ, Weiden MD. Enrichment of the lung microbiome with oral taxa is associated with lung inflammation of a Th17 phenotype. Nat Microbiol 2016; 1: 16031.

14. Hall M, Beiko RG. 16S rRNA Gene Analysis with QIIME2. Methods Mol Biol 2018; 1849: 113–129.

15. DeSantis TZ, Hugenholtz P, Larsen N, Rojas M, Brodie EL, Keller K, Huber T, Dalevi D, Hu P, Andersen GL. Greengenes, a chimera-checked 16S rRNA gene database and workbench compatible with ARB. Appl Environ Microbiol 2006; 72: 5069–5072.

16. Bisanz JE. qiime2R: Importing QIIME2 artifacts and associated data into R sessions.. 2018. Available from: https://github.com/jbisanz/qiime2R.

17. McMurdie PJ, Holmes S. Phyloseq: a bioconductor package for handling and analysis of high-throughput phylogenetic sequence data. Pac Symp Biocomput 2012: 235–246.

18. Robinson MD, McCarthy DJ, Smyth GK. edgeR: a Bioconductor package for differential expression analysis of digital gene expression data. Bioinformatics 2010; 26: 139–140.

19. McCarthy DJ, Chen Y, Smyth GK. Differential expression analysis of multifactor RNA-Seq experiments with respect to biological variation. Nucleic Acids Res 2012; 40: 4288–4297.

20. Chen Y, Lun AT, Smyth GK. From reads to genes to pathways: differential expression analysis of RNA-Seq experiments using Rsubread and the edgeR quasi-likelihood pipeline. F1000Res 2016; 5: 1438.

21. Holmes I, Harris K, Quince C. Dirichlet multinomial mixtures: generative models for microbial metagenomics. PloS one 2012; 7: e30126.

22. Davis NM, Proctor DM, Holmes SP, Relman DA, Callahan BJ. Simple statistical identification and removal of contaminant sequences in marker-gene and metagenomics data. bioRxiv 2018: 221499.

23. Keir HR, Shoemark A, Dicker AJ, Perea L, Pollock J, Giam YH, Suarez-Cuartin G, Crichton ML, Lonergan M, Oriano M, Cant E, Einarsson GG, Furrie E, Elborn JS, Fong CJ, Finch S, Rogers GB, Blasi F, Sibila O, Aliberti S, Simpson JL, Huang JTJ, Chalmers JD. Neutrophil extracellular traps, disease severity, and antibiotic response in bronchiectasis: an international, observational, multicohort study. The Lancet Respiratory medicine 2021; 9: 873–884.

24. Dicker AJ, Crichton ML, Pumphrey EG, Cassidy AJ, Suarez-Cuartin G, Sibila O, Furrie E, Fong CJ, Ibrahim W, Brady G, Einarsson GG, Elborn JS, Schembri S, Marshall SE, Palmer CNA, Chalmers JD. Neutrophil extracellular traps are associated with disease severity and microbiota diversity in patients with chronic obstructive pulmonary disease. The Journal of allergy and clinical immunology 2018; 141: 117–127.

25. Berger KI, Pradhan DR, Goldring RM, Oppenheimer BW, Rom WN, Segal LN. Distal airway dysfunction identifies pulmonary inflammation in asymptomatic smokers. ERJ Open Res 2016; 2.

26. Mac Aogain M, Narayana JK, Tiew PY, Ali N, Yong VFL, Jaggi TK, Lim AYH, Keir HR, Dicker AJ, Thng KX, Tsang A, Ivan FX, Poh ME, Oriano M, Aliberti S, Blasi F, Low TB, Ong TH, Oliver B, Giam YH, Tee A, Koh MS, Abisheganaden JA, Tsaneva-Atanasova K, Chalmers JD, Chotirmall SH. Integrative microbiomics in bronchiectasis exacerbations. Nat Med 2021; 27: 688–699.

27. Narayana JK, Aliberti S, Mac Aogáin M, Jaggi TK, Ali N, Ivan FX, Cheng HS, Yip YS, Vos MIG, Low ZS, Lee JXT, Amati F, Gramegna A, Wong SH, Sung JJY, Tan NS, Tsaneva-Atanasova K, Blasi F, Chotirmall SH. Microbial Dysregulation of the Gut-Lung Axis in Bronchiectasis. American journal of respiratory and critical care medicine 2023; 207: 908–920.

28. Mac Aogáin M, Narayana JK, Tiew PY, Ali N, Yong VFL, Jaggi TK, Lim AYH, Keir HR, Dicker AJ, Thng KX, Tsang A, Ivan FX, Poh ME, Oriano M, Aliberti S, Blasi F, Low TB, Ong TH, Oliver B, Giam YH, Tee A, Koh MS, Abisheganaden JA, Tsaneva-Atanasova K, Chalmers JD, Chotirmall SH. Integrative microbiomics in bronchiectasis exacerbations. Nature medicine 2021; 27: 688–699.

29. Mallick H, Rahnavard A, McIver LJ, Ma S, Zhang Y, Nguyen LH, Tickle TL, Weingart G, Ren B, Schwager EH, Chatterjee S, Thompson KN, Wilkinson JE, Subramanian A, Lu Y, Waldron L, Paulson JN, Franzosa EA, Bravo HC, Huttenhower C. Multivariable association discovery in population-scale meta-omics studies. PLoS Comput Biol 2021; 17: e1009442.

30. **Verma D, Stapleton M, Gadwa J, Vongtongsalee K, Schenkel AR, Chan ED, Ordway D. Mycobacterium avium Infection in a C3HeB/FeJ Mouse Model. Front Microbiol 2019; 10: 693.

31. **Andréjak C, Almeida DV, Tyagi S, Converse PJ, Ammerman NC, Grosset JH. Characterization of mouse models of Mycobacterium avium complex infection and evaluation of drug combinations. Antimicrobial agents and chemotherapy 2015; 59: 2129–2135.

32. Rogers GB, Bruce KD, Martin ML, Burr LD, Serisier DJ. The effect of long-term macrolide treatment on respiratory microbiota composition in non-cystic fibrosis bronchiectasis: an analysis from the randomised, double-blind, placebo-controlled BLESS trial. The Lancet Respiratory medicine 2014; 2: 988–996.

33. Tsay JJ, Wu BG, Sulaiman I, Gershner K, Schluger R, Li Y, Yie TA, Meyn P, Olsen E, Perez L, Franca B, Carpenito J, Iizumi T, El-Ashmawy M, Badri M, Morton JT, Shen N, He L, Michaud G, Rafeq S, Bessich JL, Smith RL, Sauthoff H, Felner K, Pillai R, Zavitsanou AM, Koralov SB, Mezzano V, Loomis CA, Moreira AL, Moore W, Tsirigos A, Heguy A, Rom WN, Sterman DH, Pass HI, Clemente JC, Li H, Bonneau R, Wong KK, Papagiannakopoulos T, Segal LN. Lower Airway Dysbiosis Affects Lung Cancer Progression. Cancer Discov 2021; 11: 293–307.

34. Wu BG, Sulaiman I, Tsay JJ, Perez L, Franca B, Li Y, Wang J, Gonzalez AN, El-Ashmawy M, Carpenito J, Olsen E, Sauthoff M, Yie K, Liu X, Shen N, Clemente JC, Kapoor B, Zangari T, Mezzano V, Loomis C, Weiden MD, Koralov SB, D’Armiento J, Ahuja SK, Wu XR, Weiser JN, Segal LN. Episodic Aspiration with Oral Commensals Induces a MyD88-dependent, Pulmonary T-Helper Cell Type 17 Response that Mitigates Susceptibility to Streptococcus pneumoniae. American journal of respiratory and critical care medicine 2021; 203: 1099–1111.

35. Gautam S, Stahl Y, Young GM, Howell R, Cohen AJ, Tsang DA, Martin T, Sharma L, Dela Cruz CS. Quantification of bronchoalveolar neutrophil extracellular traps and phagocytosis in murine pneumonia. American journal of physiology Lung cellular and molecular physiology 2020; 319: L661–l669.

36. Segata N, Izard J, Waldron L, Gevers D, Miropolsky L, Garrett WS, Huttenhower C. Metagenomic biomarker discovery and explanation. Genome Biol 2011; 12: R60.

